# Revisiting the bad luck hypothesis: Cancer risk and aging are linked to replication-driven changes to the epigenome

**DOI:** 10.1101/2022.09.14.507975

**Authors:** Christopher J. Minteer, Kyra Thrush, Peter Niimi, Joel Rozowsky, Jason Liu, Mor Frank, Thomas McCabe, Erin Hofstatter, Mariya Rozenblit, Lajos Pusztai, Kenneth Beckman, Mark Gerstein, Morgan E. Levine

## Abstract

Aging is the leading risk factor for cancer. While it’s been proposed that the age-related accumulation of somatic mutations drives this relationship, it is likely not the full story. Here, we show that both aging and cancer share a common epigenetic replication signature, which we modeled from DNA methylation data in extensively passaged immortalized human cells *in vitro* and tested on clinical tissues. This epigenetic signature of replication – termed CellDRIFT – increased with age across multiple tissues, distinguished tumor from normal tissue, and was escalated in normal breast tissue from cancer patients. Additionally, within-person tissue differences were correlated with both predicted lifetime tissue-specific stem cell divisions and tissue-specific cancer risk. Overall, our findings suggest that age-related replication drives epigenetic changes in cells, pushing them towards a more tumorigenic state.

**One sentence summary:** Cellular replication leaves an epigenetic fingerprint that may partially underly the age-associated increase in cancer risk.

## Introduction

Aging is a leading carcinogen, with cancer risk increasing over 4,000% between the ages of 25 and 65 [1]. This staggering increase is not the entire cancer narrative [2], but it is the exigent characteristic of the disease. While some cancers—like those of the bone, brain, or nervous system—are diagnosed at higher frequencies in children and adolescence [3], recent studies suggest only one-third of all cancers are linked to age-independent factors. This begs the question, to what degree is cancer preventable, as opposed to a largely unavoidable outcome of time [4, 5]? This premise, argued by Vogelstein and Tomasetti in 2015, later became known as the bad luck hypothesis [4]. The controversial theory was based on the idea that the cumulative number of divisions a cell undergoes over time is related to its propensity for tumorigenesis.

Vogelstein and Tomasetti concluded that stochastic mutation accumulation in presumed stem cells explains a far greater number of cancers than do germline predisposition and environmental factors. However, one caveat that should be considered is that their data drew from population-level statistics, rather than the individual-level. Inter-individual heterogeneity can be obscured only by measuring trends preserved across the population and tissues. Population-level statistics do measure inter-individual differences, but only differences that trend across the majority of the population. This does not exclude the likely possibility of between-person heterogeneity when considering risk of cancer within a given tissue type. The theory also relies on a presumed tissue specific stem cell as the originator of cancer, a hypothesis that has been called into question. It is an open question whether variations in rates of biological aging, interpersonal differences in cumulative cell divisions in tissues, and their accompanying molecular changes, contribute to differential risk of cancer across individuals.

Mutations are not the only – or perhaps even the most important – molecular events that result from cellular proliferation. We and others have shown that DNA methylation (DNAm) is also substantially altered as a direct function of cell division [6–9]. Further, the epigenome has been shown to undergo dramatic changes with aging and is implicated in establishing, driving and maintaining many cancers [10–13]. Coincidently, the DNA methylation changes observed in aging, cancer, and proliferation share some notable patterns. In general, they tend to be characterized by gains in methylation at promoters— especially those marked by polycomb group (PcG) factor targets—and loss of methylation in intergenic regions and repetitive elements [14]. Thus, one hypothesis is that as cells replicate in aging tissues they may also take on epigenetic signatures that are more cancer-like, making the leap to oncogenic transformation progressively more likely with time [15–17]. Further, the greater the replication rate in a given tissue, the faster this transformation may occur.

To test this, we quantified a “replication fingerprint” in DNAm data derived from extensively passaged immortalized human cells. We show this signature accumulates with aging in tissues, is stronger in tumor relative to normal tissues, appears accelerated in the normal tissue of cancer patients, and correlates with tissue-specific differences in life-time cancer risk and total stem-cell divisions.

## Results

### Identification and isolation of replication fingerprints

hTERT immortalized fetal astrocytes were serially passaged and DNAm was assessed longitudinally at each passage (Fig. 1A). We selected immortalized human fetal astrocytes as our model of choice as we reasoned that fetal cells would exhibit less fitness selection upon culturing, in comparison to adult primary cells. We also hypothesized that replication-driven DNAm changes would be cell-type independent. Additionally, immortalized astrocytes are commonly used as glial disease models due to their similarities in signaling to primary astrocytes and dramatically improved proliferation in culture, providing a physiologically relevant model with extended proliferative potential [18, 19]. The cellular lifespan of the immortalized astrocytes in our study was extended by more than 700% compared to non-immortalized astrocytes. After 73+ cumulative PDs (when we ended data collection), cells showed no sign of growth arrest, genomic instability, or telomere erosion, allowing us to better isolate the effect of replication-based epigenetic changes (Fig. 1B, Sup Fig. 1).

**Figure 1:**
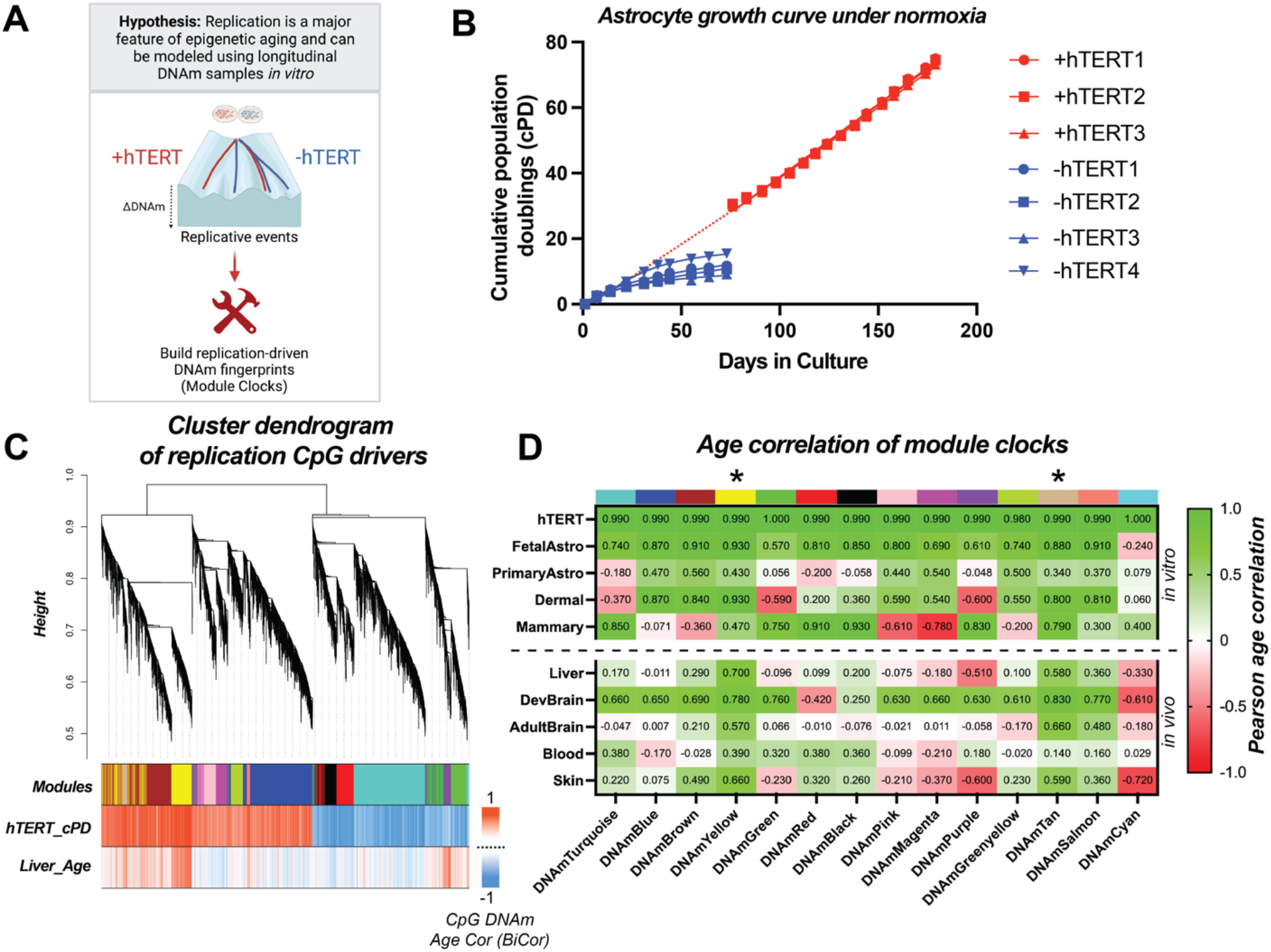
Replication-driven *epigenetic* fingerprints trained from longitudinal hTERT trajectory *in vitro* captures multi-tissue epigenetic aging. **(A)** Schematic displaying goal of cataloging epigenetic aging trajectories and signals using a longitudinal hTERT passaging model *in vitro*. **(B)** Growth curve of human astrocytes under normoxia showing growth arrest occurs after 10 passages in the absence of hTERT immortalization. **(C)** Cluster dendrogram showing the construction of the modules fingerprints via estimating the adjacencies of the input CpGs, which were the highest contributing 20,101 CpGs contributing to replication, determined from pulling the top PC loading CpGs from the hTERT trained PC measure, DNAmImmort. Briefly, adjacencies were converted to topological overlap matrices (TOMs) and then clustered based on a minimum dissimilarity score between hTERT cPD and liver aging (years) input data. hTERT_cPD and Liver_Age plot labels display the BirCor correlations (cPD or Age in years) for all 20,101 CpGs used in the clustering analysis. The goal of the clustering analysis was to produce the most physiologically relevant replication-driven aging signals. **(D)** Summary table displaying Pearson age correlation of all PC clock module measures with multi-tissue and *in vitro* validation datasets from Sup Fig. 3. Note, the correlation from the hTERT dataset used during training was excluded from the hTERT validation samples. Additionally, liver, developing brain, adult brain, blood and skin were assessed as age-correlations, fetal astrocyte and primary astrocytes were assessed as cumulative population doublings (cPD) correlations, and dermal and mammary cell lines were assessed as passage correlations due to limited cPD data information. For a complete breakdown of the age distributions for each in vivo dataset refer to Sup Fig. 3C. * = the modules (yellow/tan) that were considered the most physiologically relevant and thus were used in subsequent analysis.

As a first step towards identifying a signature of cell division, we conducted consensus network analysis to identify modules of co-methylated CpGs [20]. In brief, a subset of 20,101 CpGs were initially selected based on their normalized loading scores from elastic net selected Principal Components (PCs) that tracked with cPD in our *in vitro* model (Sup Fig. 2). Clustering of these CpGs was then carried out based on consensus clustering between our *in vitro* model and multi-age samples from human liver (N=85, 23-83 years old). This enabled us to segregate physiologically relevant replication signals from cell culture artifacts (Fig. 1C).

The association of 14 consensus modules with number of cell divisions (cPDs) *in vitro* or aging *in vivo* were assessed across a diverse spectrum of cells and tissues. Two modules—termed “yellow” and “tan”— stand out as having signatures that consistently increased with proliferation and aging (Fig. 1D, Sup Fig. 3, 4). The yellow and tan modules are the most physiologically relevant replication fingerprints, with strong correlations with age across a variety of *in vivo* tissues, including liver (r_yellow_=0.70, r_tan_=0.58, N=85, 23-83 years old), skin (r_yellow_=0.66, r_tan_=0.59, N=91, 20-90 years old), developing brain (r_yellow_=0.78, r_tan_=0.83, N=173, 0-18 years old), and adult brain (r_yellow_=0.57, r_tan_=0.66, N=502, 18-97 years old), and blood (r_yellow_=0.39, r_tan_=0.14, N=2478, 40-92 years old) (Fig. 1D, Sup Fig. 3C). They also tracked serial passaging of primary cultures of fetal astrocytes (r_yellow_=0.93, r_tan_=0.88), primary astrocytes (r_yellow_=0.43, r_tan_=0.34), dermal fibroblasts (r_yellow_=0.93, r_tan_=0.8) and mammary fibroblasts (r_yellow_=0.47, r_tan_=0.79) (Fig. 1D). For these reasons, the CpGs in the yellow and tan modules served as the basis for all other downstream analyses in the study. Further details on the training and validation of these CpG modules is found in the computational section of the Methods.

### Replication fingerprints are enriched in PRC2, pluripotency factors, and cell cycle regulator domains

Overall, the yellow and tan module CpGs were equally distributed across the genome and were not located in any one region (Sup Fig. 2). They were, however, highly enriched in PRC2, pluripotency factors and cell cycle regulator domains. Enrichment was assessed using the Cistrome database, which tests overlap for specific genomic locations of interest based on biorepository data from ENCODE and includes information on specific histone marks, transcription factors (TF), and chromatin regulators (Sup Fig. 5). Both modules were enriched in regions marked by H3K27me3, which included both somatic and stem cell datasets (Fig. 2A). When testing for overlap with known TF binding sites, we observed enrichment in the yellow module with TFs that formed an interactive STRING protein network with PRC2 domains, such as the catalytic subunit EZH2 and interacting cofactors SUZ12, EP300, JARID2 and TRIM28 (Fig. 2B). PRC2 elements were also some of the most enriched by absolute score (Sup Fig. 5). This is of particular interest since PRC2 domains are implicated in development and maintenance of many cancer types [21]. PRC2 is a trimeric multiprotein complex (EZH2/EED/SUZ12), although it interacts with many other upstream and downstream co-factors like EP300, JARID2 and TRIM28, all of which have profound impacts on controlling cellular differentiation, signaling and genome-wide regulation [22]. Our enrichment data suggests the epigenome may contribute to replication-driven dysregulation through many PRC2 component interactions, including both upstream and downstream regulators in addition to the trimeric core. CpGs in the tan module exhibited enrichment for additional noteworthy TFs, including KLF4, RAD51 and STAT3 (Fig. 2B). KLF4 is a pluripotency factor and considered to be a tumor suppressor in many types of cancer, while RAD51 and STAT3 may pre-dispose cells to cancer through faulty DNA repair or signal activation [23–30]. Importantly, the yellow and tan module TF enriched hits were distinct, with the only overlap being CTCF, suggesting they represent two independent replication signals.

**Figure 2:**
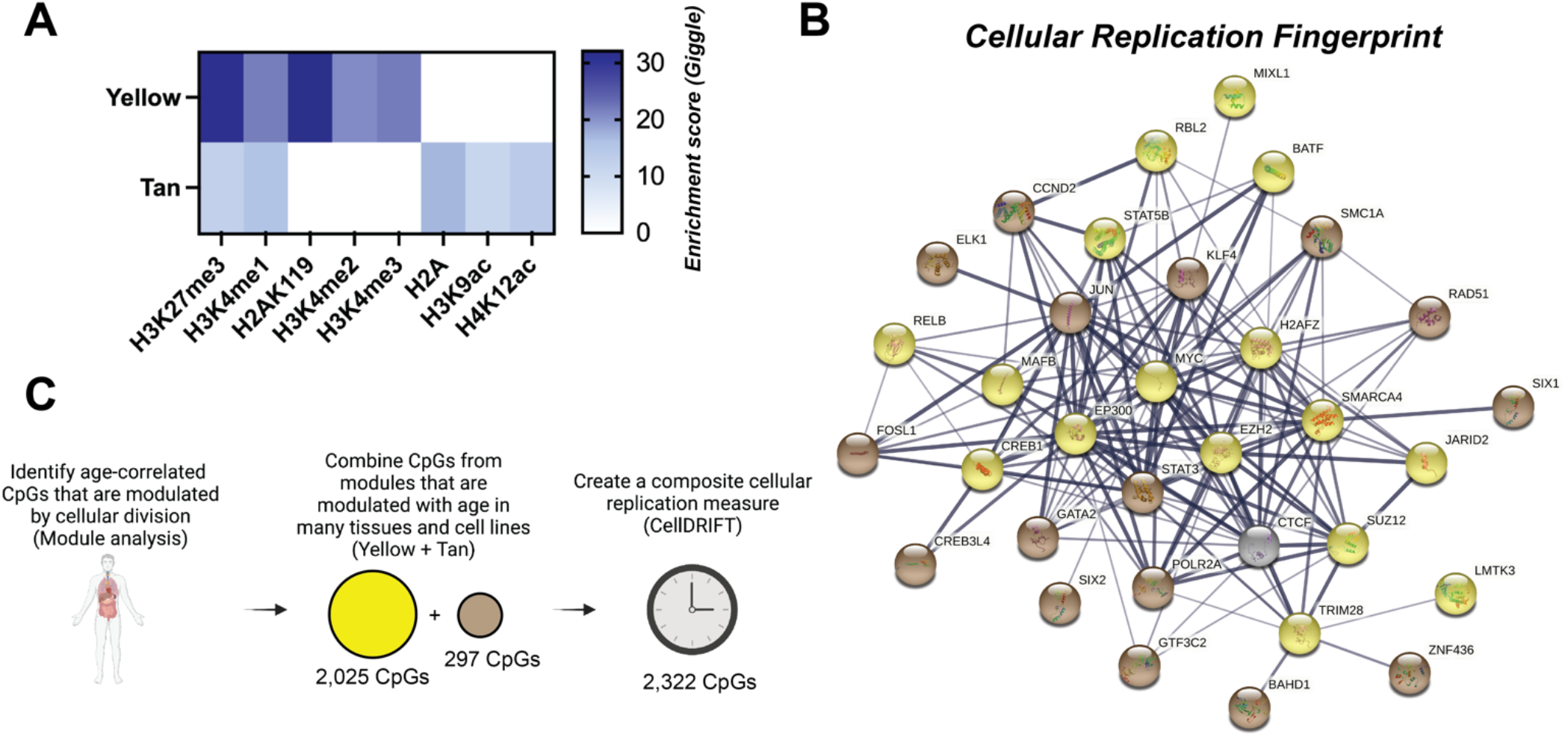
Replication-driven aging signals are enriched in Polycomb group factor 2 domains and cell cycle regulators. **(A)** Heatmap demonstrating histone enrichment score of module CpGs from Cistrome genome enrichment analysis. **(B)** String analysis of transcription factors and chromatin regulators enriched in CpGs from Yellow and Tan modules. Note, histone enrichment (A) and transcription factor/chromatin regulator fingerprint analysis (B) was determined from conducting Cistrome genome enrichment analysis on the top 100 kME CpGs from each module. The top 20 enriched genes were analyzed in the string analysis (B), with weakly connected nodes excluded. Final network strength and connectivity is based on line thickness and color grouping is based on module occupancy, with yellow = yellow module, tan = tan module and grey = both modules. See Supplemental Figure 5 for additional Cistrome genome enrichment analysis. **(C)** Schematic displaying selection process of isolating CpGs that are established by repeated cellular divisions and are also modulated with age in many tissues (yellow + tan module CpGs). The composite CpGs were used to create the measure CellDRIFT, which is used in subsequent analysis.

### Evaluation of replication fingerprint (CellDRIFT) in cancer patients, high caner-risk individuals, and tissues with varying replicative history

To directly test the associations between these signals and changes in aging and cancer, we created a composite measure from the 2,322 CpGs in the yellow and tan modules, termed CellDRIFT (**<u>Cell</u>**ular **<u>D</u>**ivision and **<u>R</u>**eplication **<u>I</u>**nduced **<u>F</u>**ingerprin**<u>T</u>**) (Fig. 2C, Sup Fig. 6, Data Fig. 1).

Malignant cells outgrow healthy cells via a number of proliferative mechanisms, including reduced activity of tumor suppressor genes, increased expression of oncogenes, chromatin dysregulation and altered transcription [14, 31]. Many of these features, particularly epigenetic remodeling, may be acquired progressively over time, prior to transformation. Unlike the bad luck and/or two-hit hypotheses, which describe mutations or a “hit” as the cancer prone tipping point, the gradual and subversive epigenetic remodeling that likely is occurring as cells “tick” throughout life provides a path for understanding aging and cancer risk years prior to phenotypic penetrance. While our measure was not trained on cancer in any way, we hypothesize that CellDRIFT will capture aspects of premalignant changes as they accumulate and we predict that 1) Cell DRIFT increases in tumors compared to normal tissues, 2) will be higher in normal tissues from individuals who develop cancer versus healthy controls, and 3) will be higher in tissues with greater proliferative activity and subsequent cancer risk (Fig. 3A).

**Figure 3:**
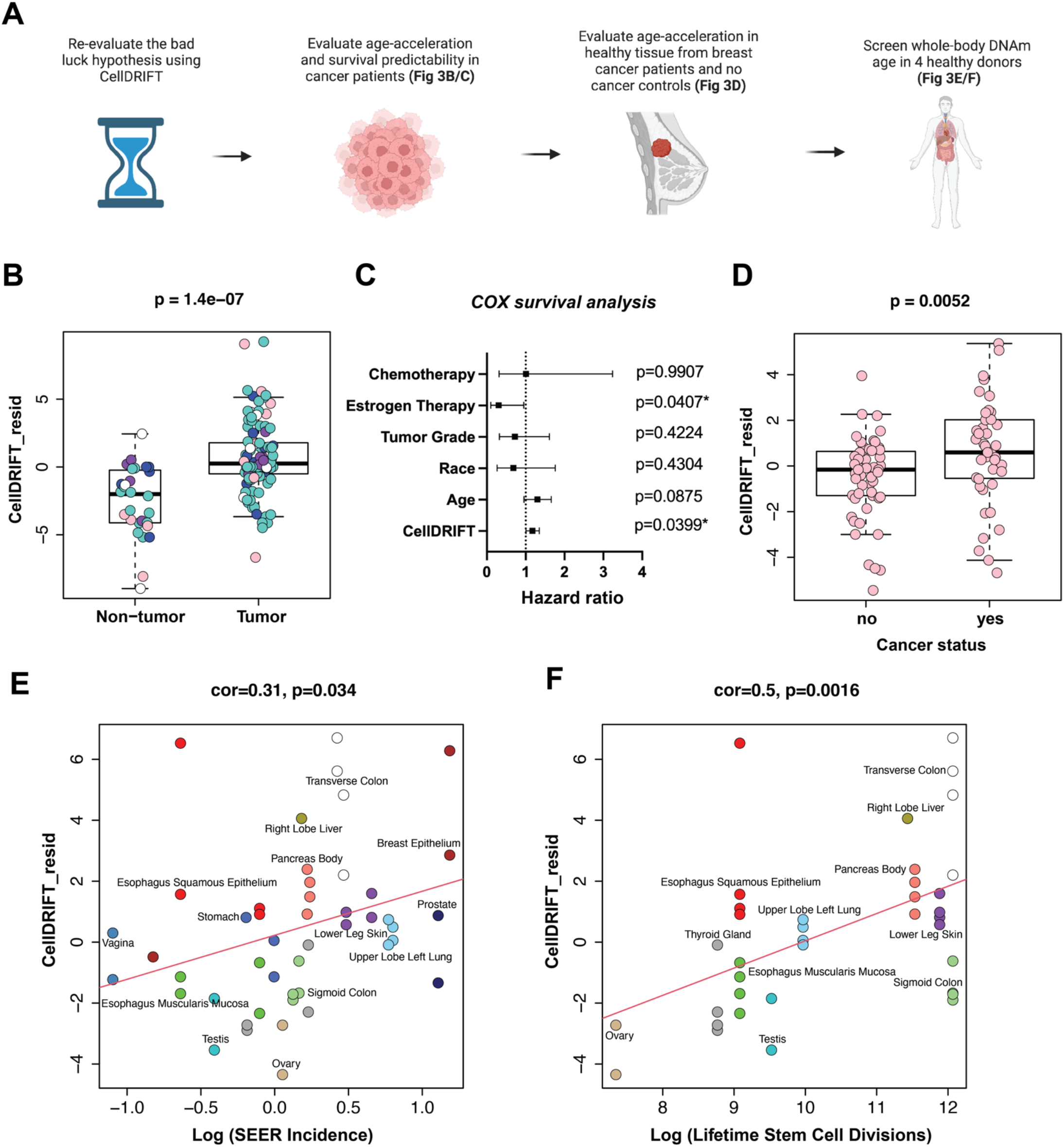
CellDRIFT predicts accelerated epigenetic risk in cancer tissue and healthy tissue of breast cancer patients, and is correlated with cancer risk and life-time stem cell divisions in whole-body tissue analysis. **(A)** Schematic displaying process of investigating the hypothesis that cancer patients or individuals with accelerated epigenetic aging may possess DNAm signals that may put these individuals at an increased risk for diseases like cancer. **(B)** Pooled cancer and normal tissue from breast, colon, lung, pancreas and thyroid cancer patients and controls, evaluated via CellDRIFT for DNAm acceleration in cancer tissue (GSE53051). Teal=Thyroid, Pink=Breast, White=Lung, Purple=Pancreas and Blue=Colon. DNAmAge scores were residualized by age, sex and tissue type. **(C)** COX survival analysis of breast cancer patients from GSE37754, demonstrating CellDRIFT is predictive of patient survival. Note, the hazard ratio for Age and CellDRIFT was calculated as a risk increase per 10 years of life. **(D)** Differences in CellDRIFT epigenetic risk in healthy breast tissue of known breast cancer patients and participants with no history of prior breast cancer (Rozenblit et. al 2022) [37]. DNAmAge scores were residualized by age prior to analysis. **(E)** Whole-body DNAmAge analysis from CellDRIFT in relation to propensity for cancer (SEER incidence per 100,000 people) of 14 different tissues from 4 individuals (ENTEx study). **(F)** Plot showing correlation between lifetime stem cell divisions and DNAmAge from CellDRIFT in non-zero cancer risk tissues from C. Note, only tissues with reported lifetimes stem cell division from Volegstein et. al 2015 were analyzed. See Supplemental Figure 7 for a complete analysis of all 29 tissues, inclusive of zero risk cancer tissues. DNAmAge scores were residualized by age prior to analysis.

To test this, we first analyzed CellDRIFT in cancers of the thyroid, breast, lung, pancreas and colon. We observed a significant increase in CellDRIFT (adjusted for age and tissue type) among the pooled tumor samples in comparison to corresponding normal tissues (p=1.4e-0.7) (Fig. 3B). We followed up by analyzing a larger breast cancer cohort with reported survival data and similarly found that CellDRIFT from tumors is predictive of overall survival, even after adjusting for age, treatment, race, and tumor grade (p=0.0399) (Fig. 3C). This suggests that the epigenetic changes modeled by CellDRIFT capture cancer aggression and fitness and may even be useful for cancer pathologists in the future as a secondary diagnostics and prognostic endpoint.

The next question, which we proposed would be the most important, but also the most difficult to detect, was determining if CellDRIFT could predict accelerated drift–or high-risk patients–in pre-diseased healthy tissues. To accomplish this, we evaluated CellDRIFT in normal breast tissue from patients with and without diagnosis of breast cancer (prior to treatment), hypothesizing that individuals who develop cancer in a particular tissue may do so as a result of more advanced and deleterious epigenetic modifications in the normal aging tissue prior to tumorigenesis – as captured by CellDRIFT (Fig. 3D). As hypothesized, we find CellDRIFT is elevated in normal tissue from breast cancer patients compared to individuals who never had breast cancer (p=0.0052, Fig. 3D). Based on the results, the conclusion stands that individual differences may exists that precede a stochastic occurrence of a “bad luck” event that drives cancer formation.

Finally, in accordance with the findings from Vogelstein and Tomasetti, we reasoned that not all tissues would display the same replicative DNAm signatures, and that cancer susceptibility (lifetime tissue-specific cancer risk) and tissue-specific stem cell division rates (replicative activity) would correlate with the degree of epigenetic changes captured by CellDRIFT in various tissues. To test this, we estimated CellDRIFT in samples from ENTEx, which profiled 29 tissues from 4 patient donors [32]. For our primary analysis, we restricted tissues to those with reported lifetime cancer risk (according to the NCIs Surveillance, Epidemiology, and End Results (SEER) program). Our results showed that CellDRIFT was positively correlated with both tissue-specific cancer risk (cor=0.31, p=0.034) (Fig. 3E) and lifetime stem-cell divisions (cor=0.5, p=0.0016) (Fig. 3F). Overall, this suggests that more proliferative tissues may have greater CellDRIFT and this may explain the higher propensity for cancer over the lifetime of that tissue. It is also noteworthy that the stem-cell division prediction traversed nearly 6 orders of magnitude (i.e ovary vs. transverse colon), demonstrating the tight link between replicative activity and epigenetic regulation. Additionally, even when near zero cancer risk tissues, like the ascending aorta and gastrocnemius medialis, were included as a sensitivity analysis, the association holds (cor=0.3, p=0.0031) (Sup Fig. 7), suggesting low-cancer risk tissues exhibit less epigenetic drift potentially because of their low replicative activity. In short, CellDRIFT provides a tool for future studies of replication-associated epigenetic mechanisms that may underly Vogelstein and Tomasetti’s initial observation.

### Transient re-setting of replication fingerprints occurs with OSKM re-programming

Our final question was to determine if cellular re-programming [30] can re-set or modulate CellDRIFT. We analyzed time-course data from human fibroblasts reprogrammed to iPSCs and tracked CellDRIFT throughout the three phases of Yamanaka factor reprogramming: initiation, maturation, and stabilization. No change was seen during the early initiation phase of reprogramming. However, we observed a dramatic decrease in the CellDRIFT signal during the maturation phase, which coincides with dedifferentiation or transition to pluripotency (cor=−0.9, p=0.00039) (Fig. 4A). Upon passaging in the stabilization phase, we observed a slight “uptick” in CellDRIFT, though it did not reach statistical significance, potentially due to a lack of statistical power (cor=0.58, p=0.10) (Fig. 4A). For this reason, we analyzed an additional iPSC and ESC dataset that reported extended passaging (Fig. 4B). In both the dermal fibroblast derived iPSC and ESC cell lines, passaging strongly induced further CellDRIFT (cor=0.74, p=5.4e-5; cor=0.77, p=0.00012, respectively), suggesting pluripotent cells, despite possessing long-term passaging abilities [33, 34], are not immune to replication related epigenetic drift (Fig. 4B).

**Figure 4:**
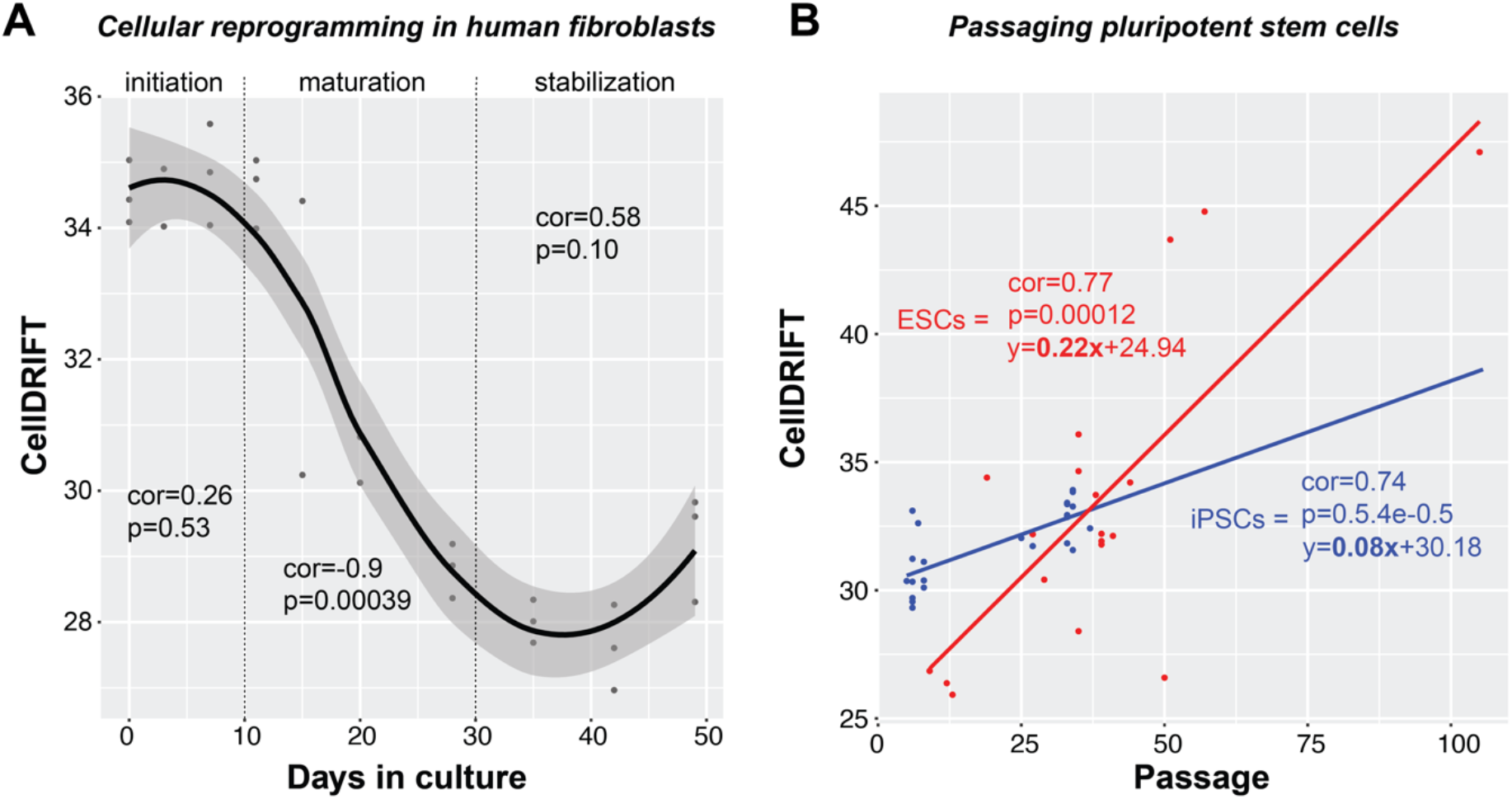
CellDRIFT is turned-back upon reprogramming in human fibroblasts, but is not stabilized with additional passages. **(A)** Scatterplot showing reprogramming trajectory in CellDRIFT from OSKM re-programmed human dermal fibroblasts from GSE54848, analyzed separately for initiation, maturation and stabilization phases. **(B)** Scatterplot demonstrating passaging increases CellDRIFT in pooled iPSC cells re-programmed from human dermal fibroblast’s via OSKM and embryonic stem cells (ESCs). Note, 11 different iPSC cells lines were included, which were generated from 11 clones and 15 ESCs cell lines were included (GSE31848). Correlation and statistical significance were determined via Pearson correlations in each phase and in the extended passaging plot.

#### Interpretations

Our study defines a signature of epigenetic remodeling observed *in vitro* as a “pure” function of DNA replication. We show that this signature increases as a function of age *in vivo*, and provides evidence to suggest it may underly an age-related transition of tissues from a (youthful) state of normal tissue and cell functioning towards an (aged) tippingpoint of tumorigenic transformation. The epigenetic changes captured in our model are consistent with prior characterizations in aging and cancer, implicating EZH2 binding sites [35] and chromatin accessibility hotspots [36]. The key takeaway is that by using a signature of molecular changes arising via replication (presented here as CellDRIFT), it is possible to quantify and track the inevitable entropic disorder that occurs with aging. While tissue-specific stem cell division rates may set a baseline risk of cancer development in various tissues, individual differences in cancer risk may not be entirely luck based and instead might in-part reflect differential epigenetic aging rates. As such, there is a need for research into interventions to slow or reverse the accumulation of epigenetic changes with age. While applications like epigenetic reprogramming demonstrate modulation is possible, issues surrounding dedifferentiation of cells, as well as feasibility and targeted efficiency mean such techniques are long from practical applications. This begs the question of whether positive lifestyle factors may modulate our luck beyond chance.

## Supporting information

Module_CpGs

## Supplemental Figures

**Supplemental Figure 1:**
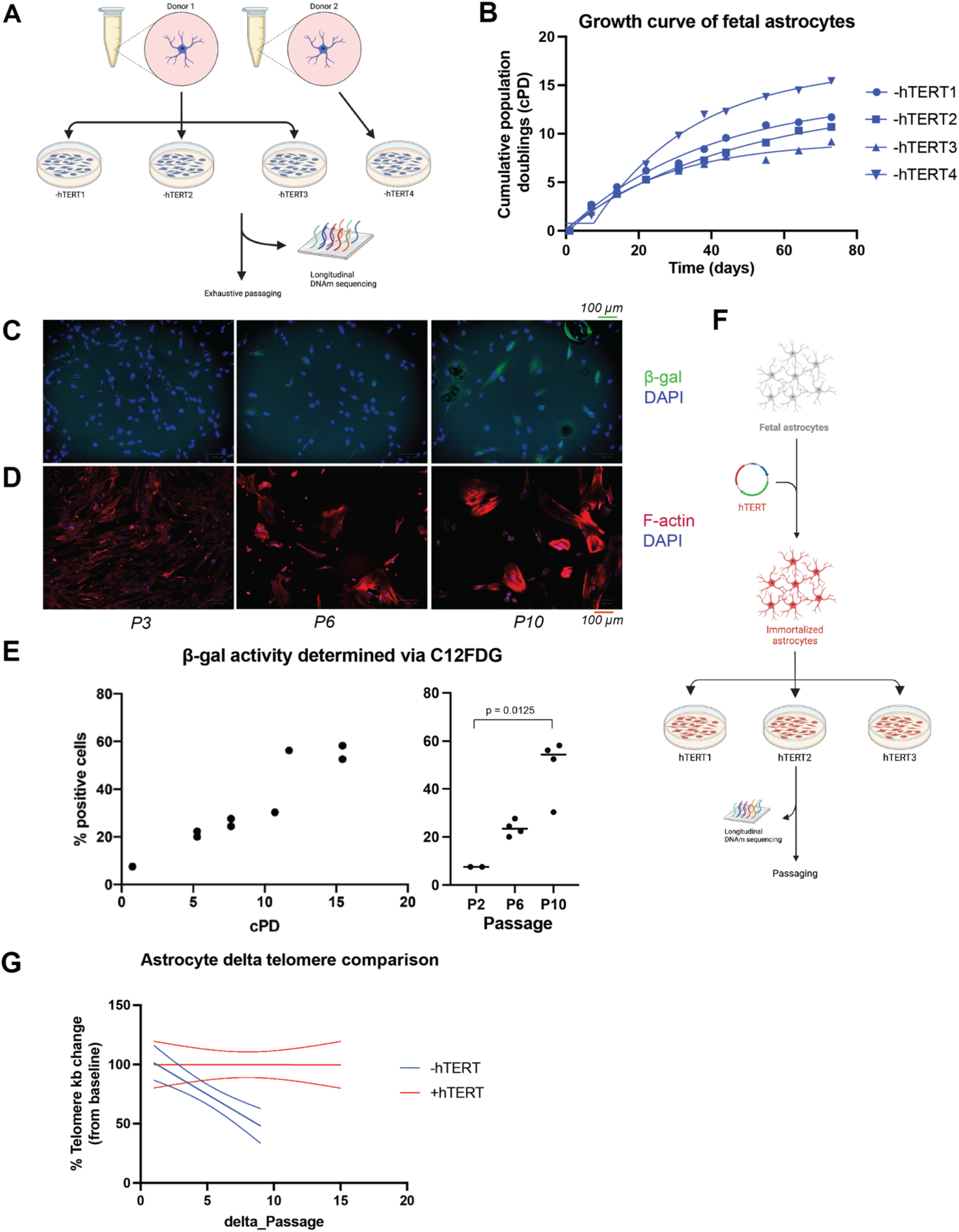
Astrocyte model development and passaging. **(A)** Schematic displaying method of creating the 4 mortal-fetal astrocyte cell lines (- hTERT1-4). 2 donors were used (Donor 1 = -hTERT1-3 and Donor 2 = -hTERT4). All replicates were then exhaustively passaged until senescence was achieved. **(B)** Plot displaying growth rate of mortal astrocytes, where growth arrest and senescence was achieved after 10x passages. **(C-D)** Representative confocal microscopy images of mortal-fetal astrocytes at P3, P6 and P10, displaying increase in senescence (β-gal) and enlarged cellular morphology (F-actin), counterstained against DAPI. **(E)** Image-J quantified β-gal activity of confocal microscopy images. **(F)** Schematic displaying method for creating the 3 immortalized-fetal astrocyte cell lines (+hTERT1-3). Note, 1 donor was used and following successful immortalization, the cells were split into 3 different cell lines (+hTERT1-3), which were then extensively passaged. Note, at the time of stopping the experiment the cells were P27, with no signs of growth arrest or senescence. Longitudinal DNAm from P13-P27 were used in subsequent PC clock creation of DNAmImmort, module clock creation and CellDRIFT. +hTERT1-2 were used in training and +hTERT3 was used as validation. **(G)** Absolute telomere length assessment of mortal (-hTERT) and immortalized (+hTERT) astrocytes demonstrating telomere erosion occurs in the absence of hTERT immortalization.

**Supplemental Figure 2:**
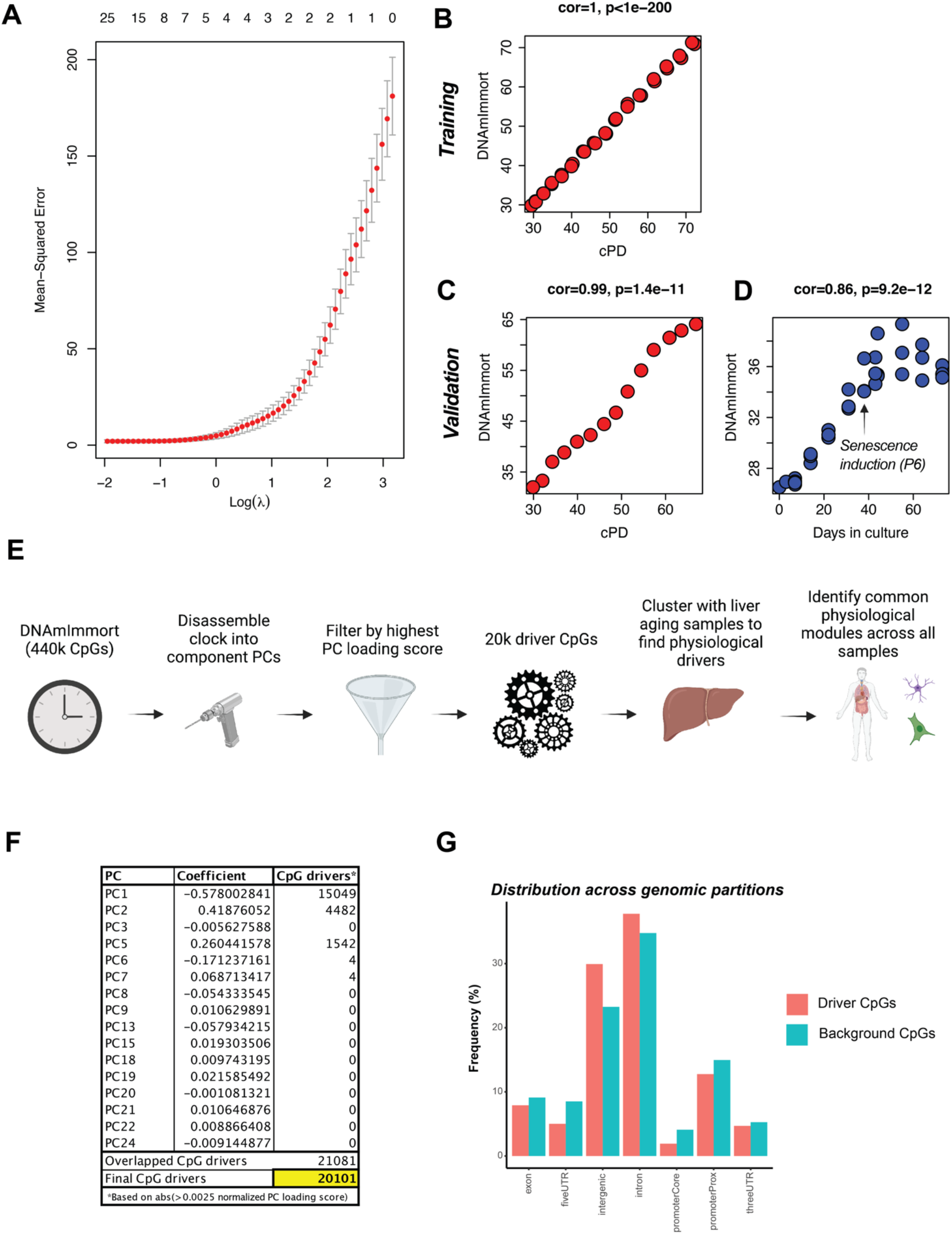
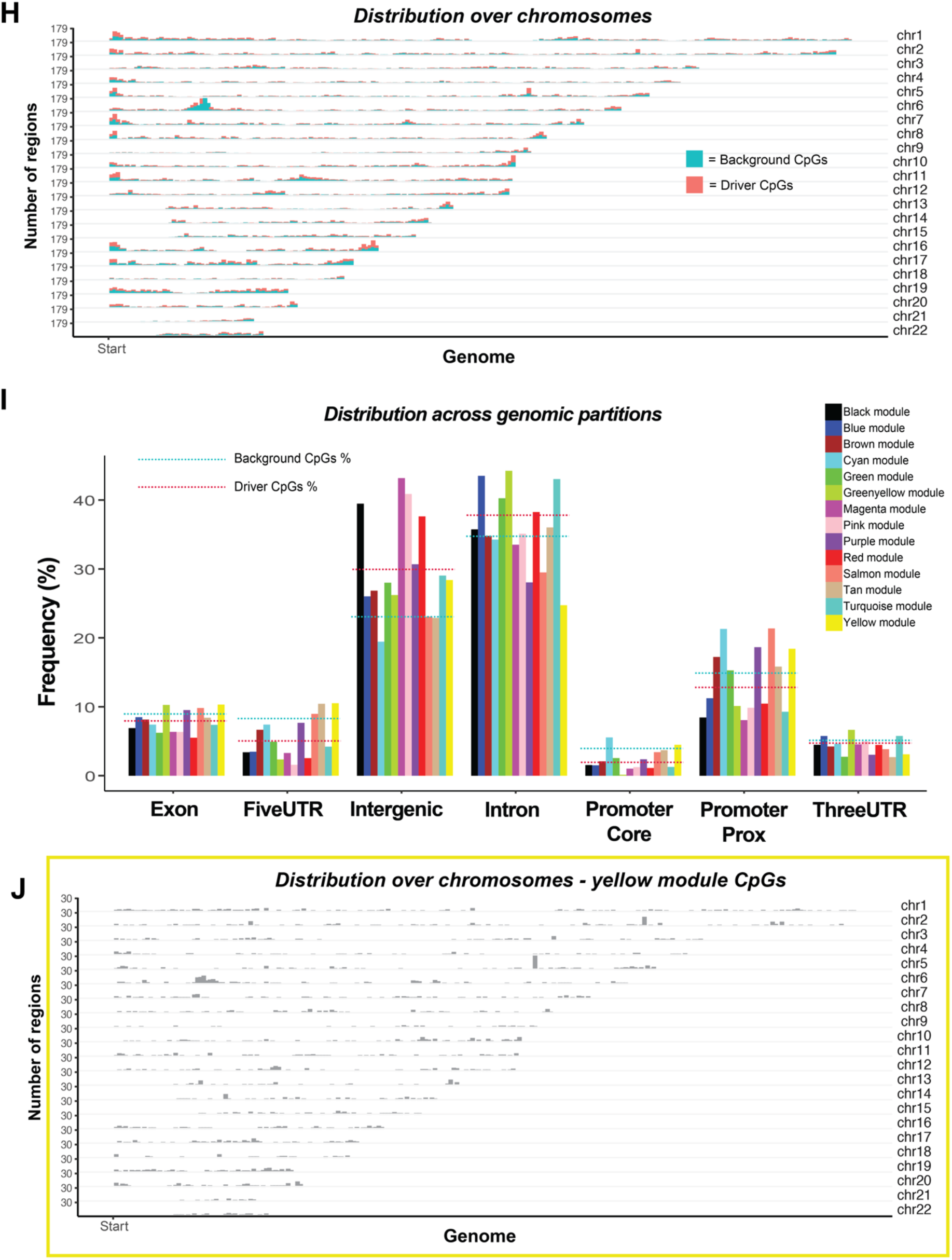
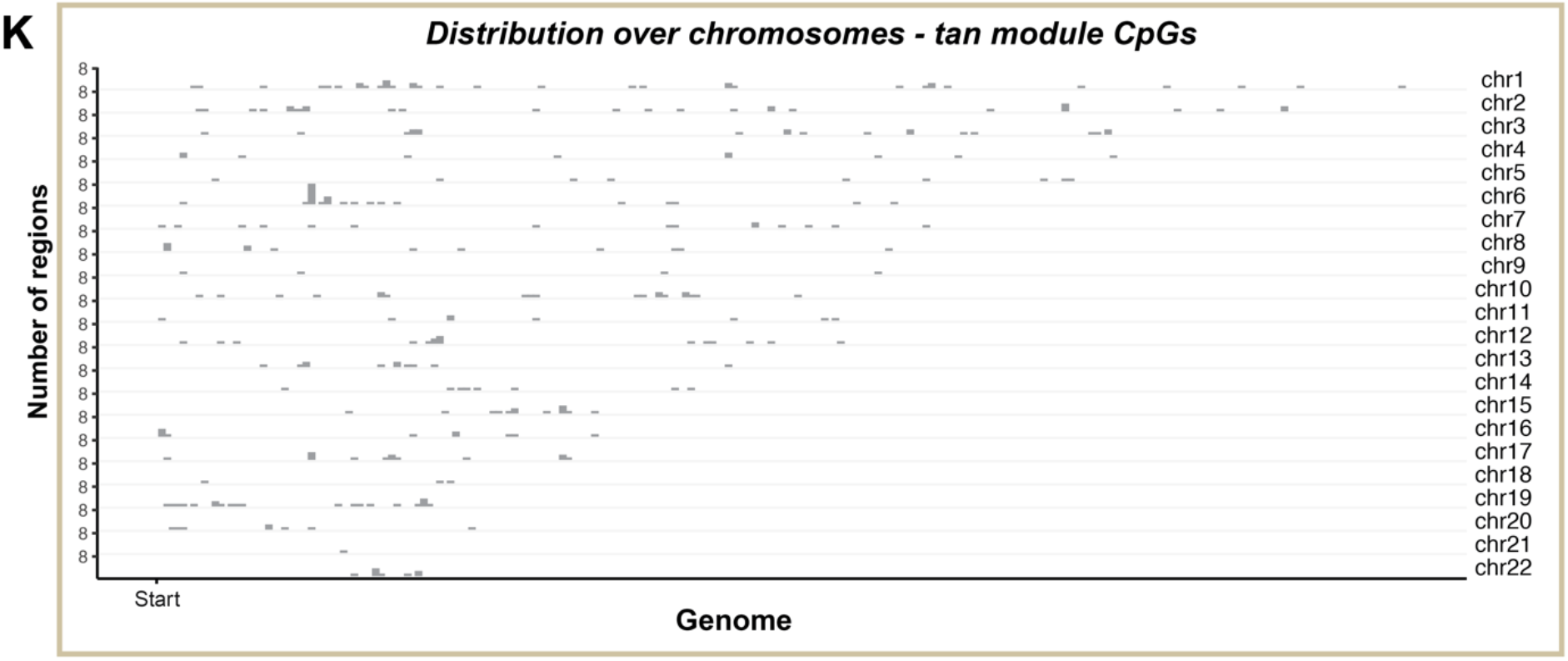
Extracting cellular replication CpG drivers. **(A)** Elastic net lambda minimum selection of PCs for inclusion in final model, DNAmImmort. **(B)** hTERT PC clock – DNAmImmort – trained from hTERT1 and hTERT2 replicates and using cPD as input variable. **(C)** DNAmImmort validation data using hTERT3 replicate not used in model training and **(D)** assessment of DNAmImmort measure in mortal (-hTERT1-3) astrocytes, displaying reduced DNAm rate upon senescence induction. **(E)** Schematic displaying workflow of breaking the DNAmImmort measure into module drivers by conducting consensus clustering on the 20,101 CpG drivers, as determined by normalizing the PC loading scores. Note, hTERT1-2 replicates and liver aging samples were used in the clustering analysis to produce the 14 modules, then using the new module CpGs module clocks were trained using the same hTERT1-2 replicates and cPD as the input variable. **(F)** Summary table of all module components in PC clock, DNAmImmort, with extracted driver CpGs (20,101) selected by pulling CpGs with a normalized (absolute value of elastic net coefficient) PC loading score of >0.0025. In the selection analysis each PC was analyzed independently. **(G)** Genomic distribution plot displaying CpG locations/regions of driver and background CpGs, showing enrichment in intronic and intergenic regions of the DNAmImmort driver CpGs. **(H)** Chromosome distribution plot generated by LolaWeb displaying 20,101 driver CpGs vs. random 20,101 background CpGs from the original 440k sex-excluded CpGs of the original training dataset. **(I)** Genomic partition distribution across all module CpGs presented in Fig. 1C, plotted using LolaWeb. Dotted lines represent the genomic partition frequency of the 20,101 driver CpGs and randomly sampled 440k background CpGs. **(J)** Chromosome distribution plot generated by LolaWeb displaying yellow module CpGs. **(K)** Chromosome distribution plot generated by LolaWeb displaying tan module CpGs.

**Supplemental Figure 3:**
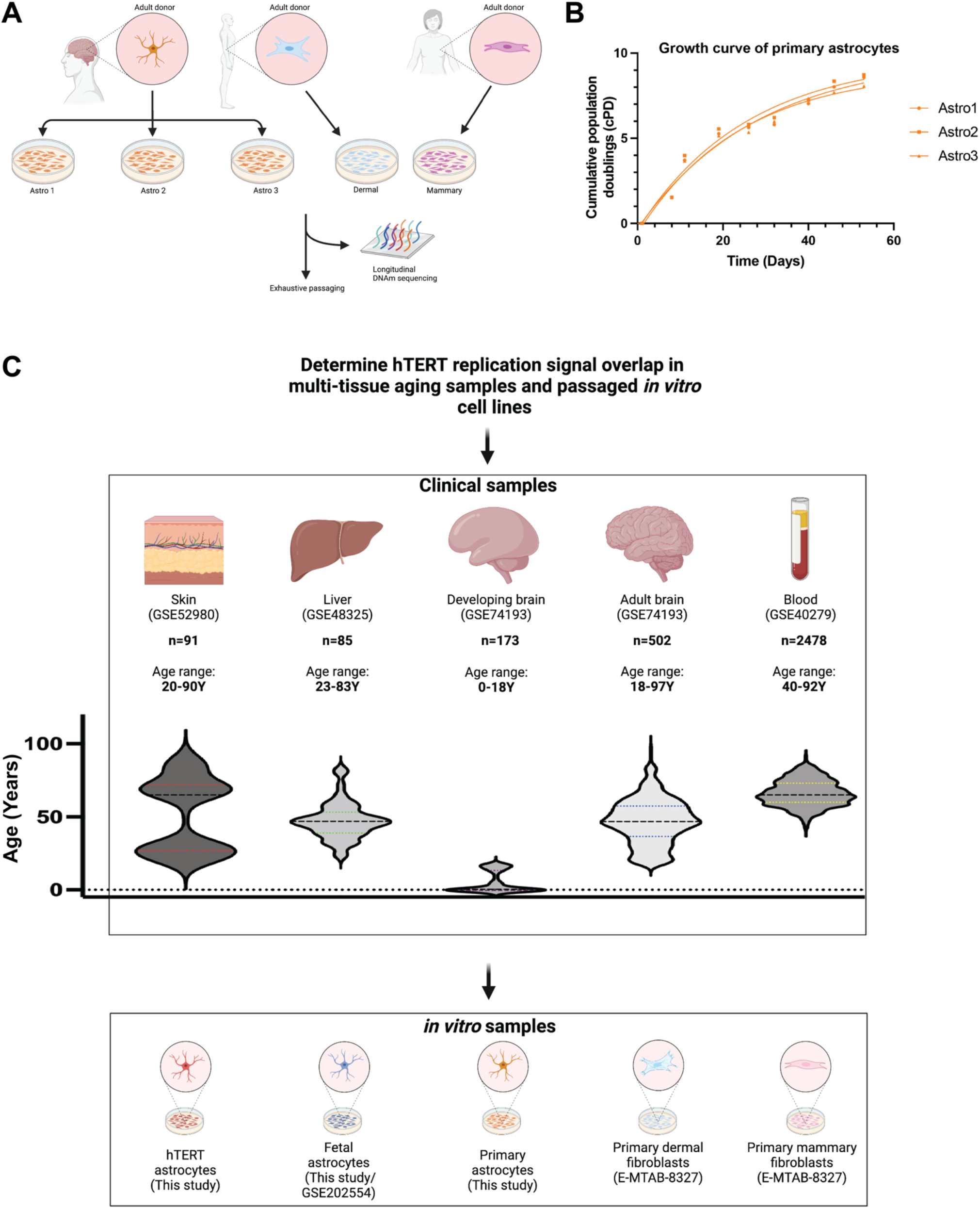
Clustering input and validation data for determining module drivers. **(A)** Schematic displaying method of extracting the primary cell lines used in the validation analysis of this study. Note, primary dermal and mammary fibroblasts were extracted by the authors of E-MTAB-8327. **(B)** Plot displaying the cumulative population doublings of primary astrocytes exhaustively passaged 10x, which was also when growth arrest was achieved. **(C)** Summary schematic of all multitissue clinical data and *in vitro* data used in the training and validation of module clock measures and CellDRIFT. Note, population distribution of all clinical datasets are displayed with violin plots.

**Supplemental Figure 4:**
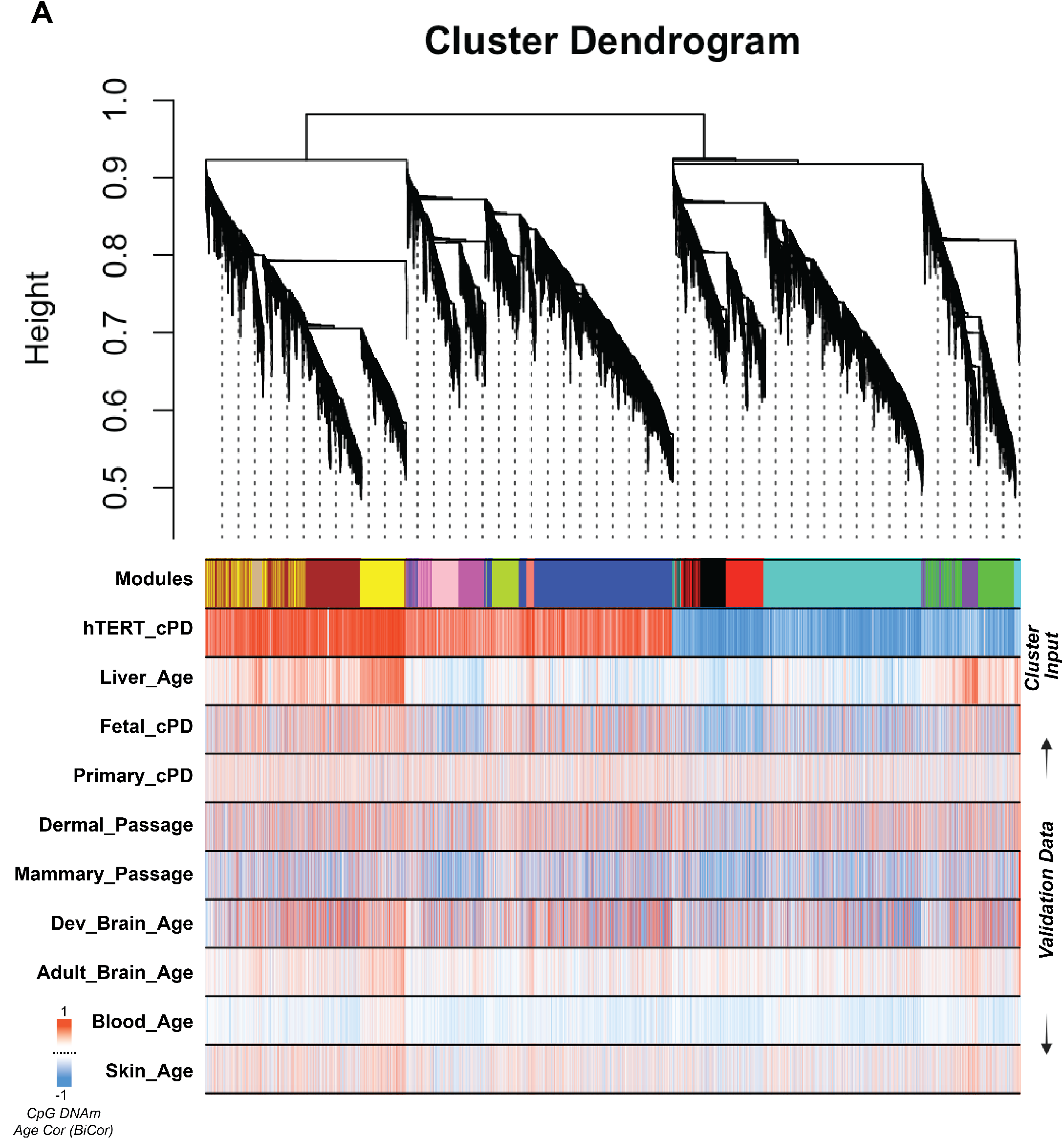
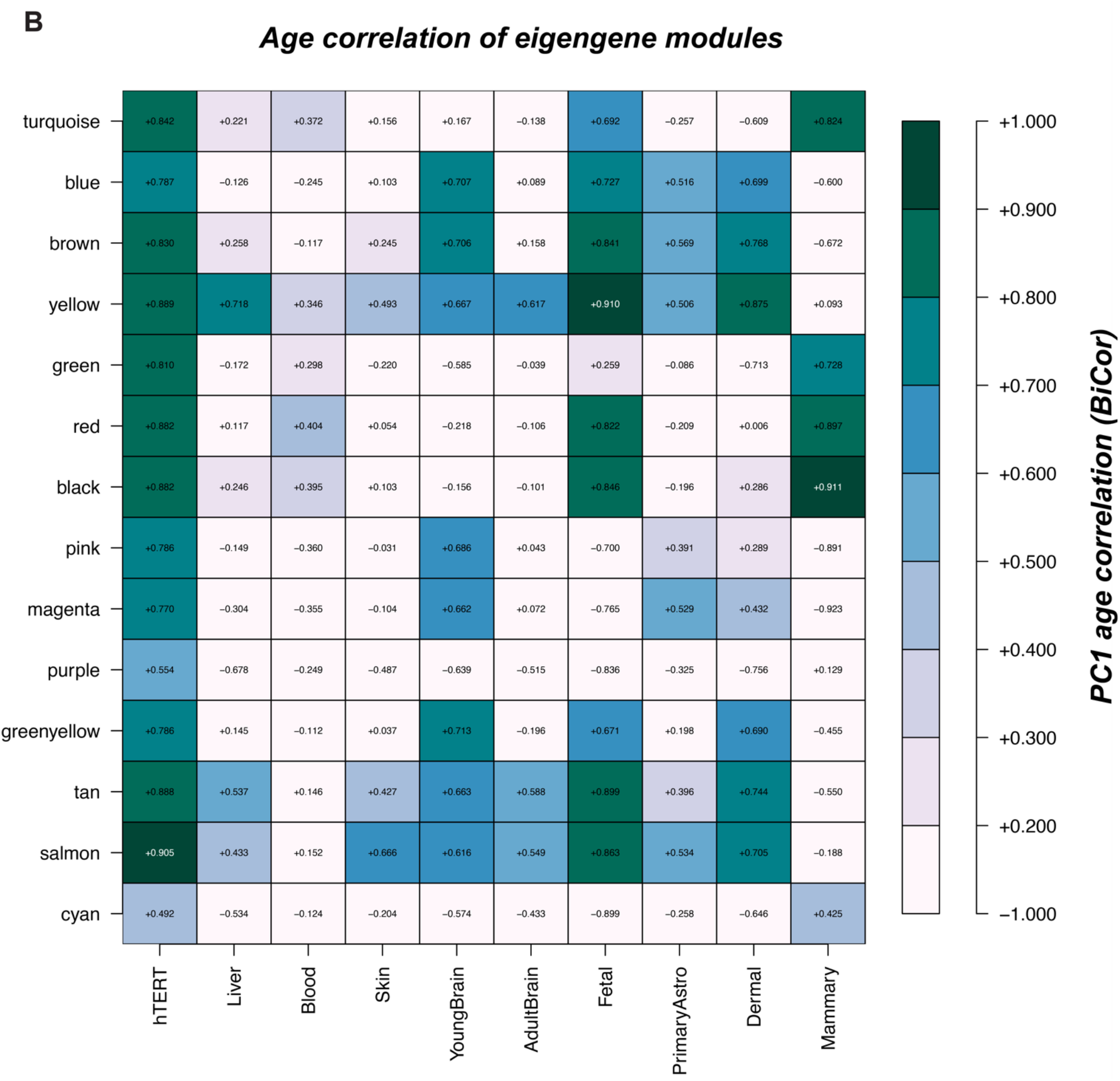
Multi-tissue and *in vitro* assessments of module relevance. **(A)** Cluster dendrogram with all input (hTERT + liver aging) and validation *in vitro* and *in vivo* samples. Plot labels display the BirCor correlations (cPD, Age in years or Passage) for all 20,101 CpGs used in the clustering analysis. **(B)** Eigengene age correlation of module CpGs in multi-tissue and *in vitro* validation datasets. PC1 validation of module CpGs analyzed in relation to hTERT validation directionality.

**Supplemental Figure 5:**
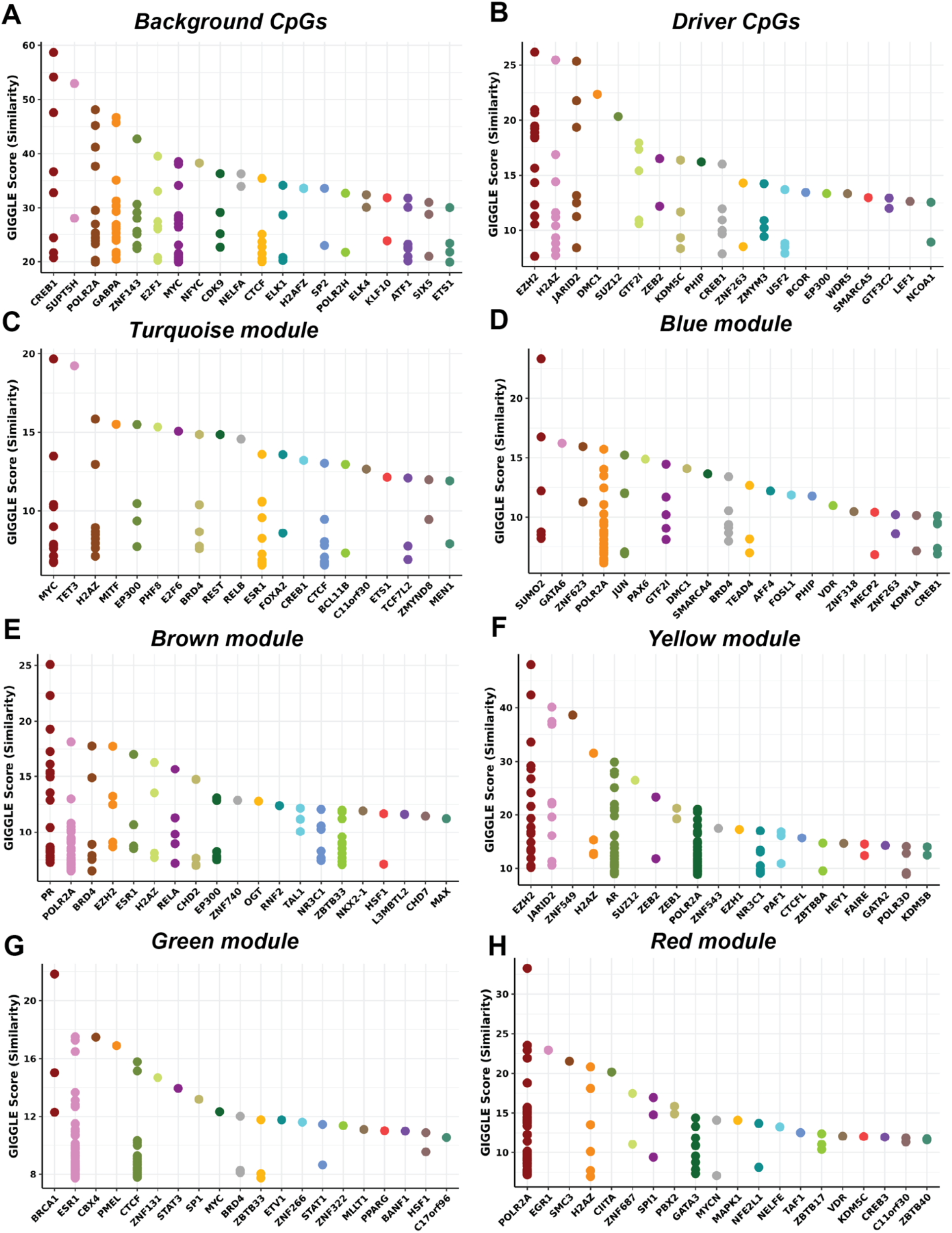
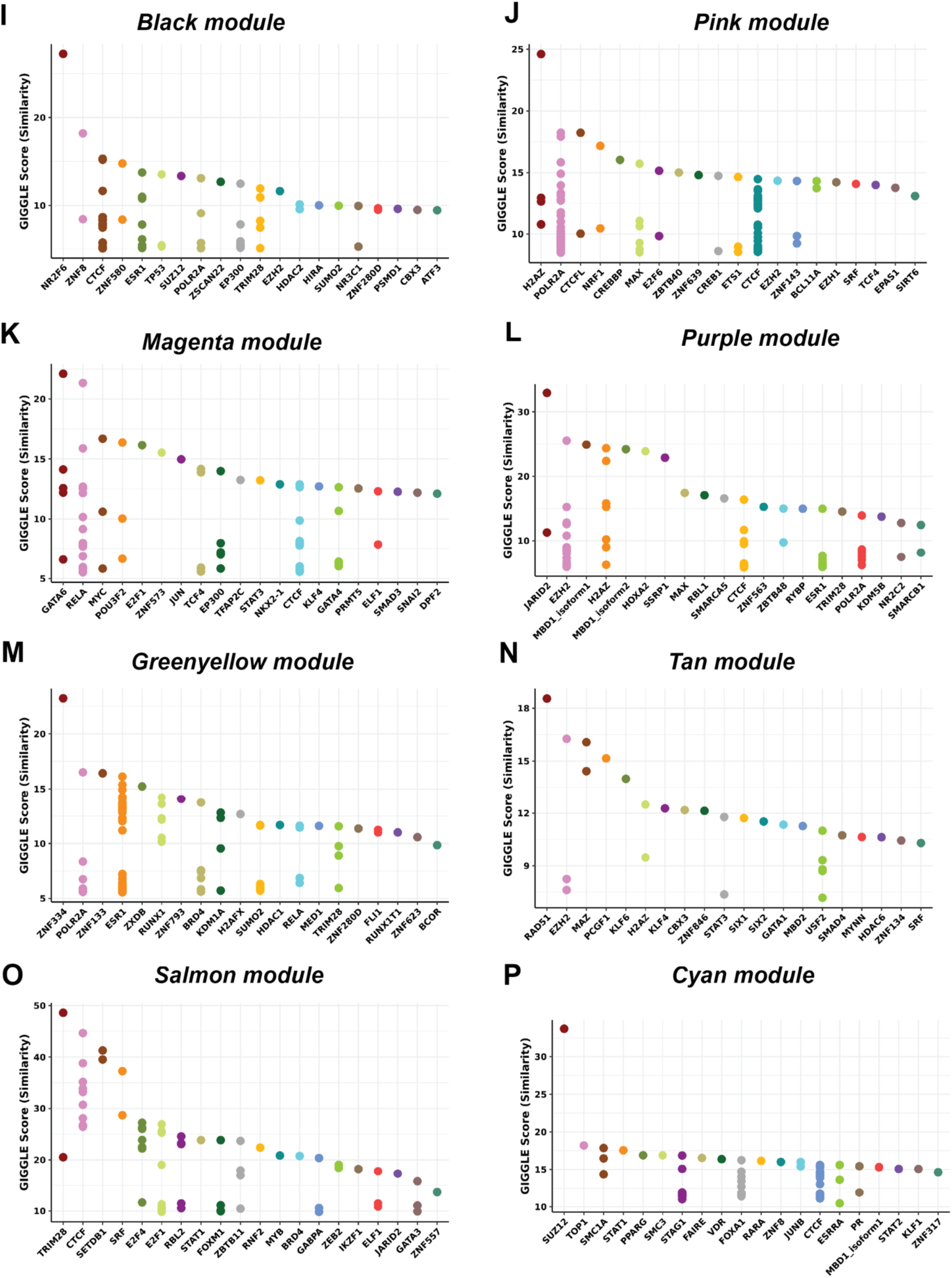
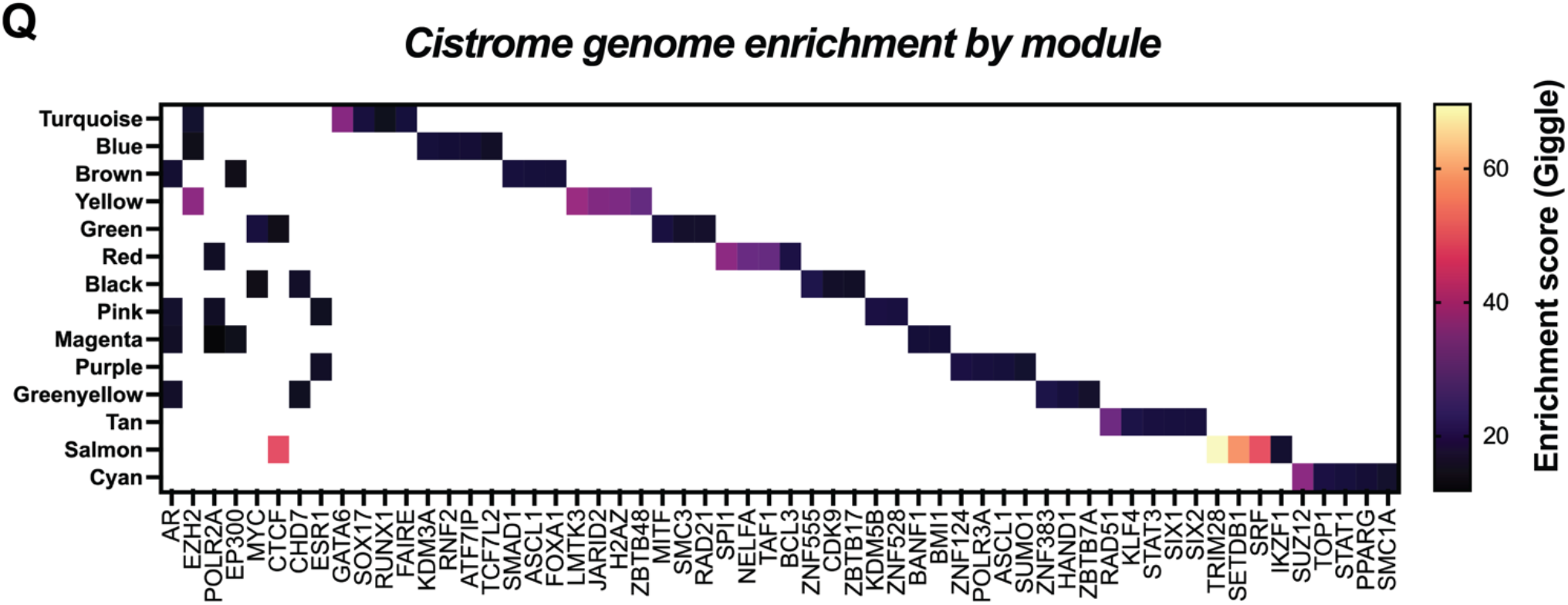
Cistrome analysis of CpGs from each module. **(A-P)** Cistrome genome enrichment plots displaying known chromatin regulators and transcription factor sites that interact with each module. All CpGs from each module were included in the analysis. **(Q)** Summary plot of top 5 enriched genes for each module. Note, the summary module enrichment analysis used the top 100 CpGs from each module, determine from the kME score. Enriched genes were normalized by selecting 100 background CpGs from the original 440k training dataset and correcting for each GSM_IDs Giggle score. Enrichment analysis is displaying the average Giggle score across all GSM_IDs, with the top 5 for each module plotted. Giggle score is a rank of genome significance between genomic locations of query file and thousands of genome files from databases like ENCODE.

**Supplemental Figure 6:**
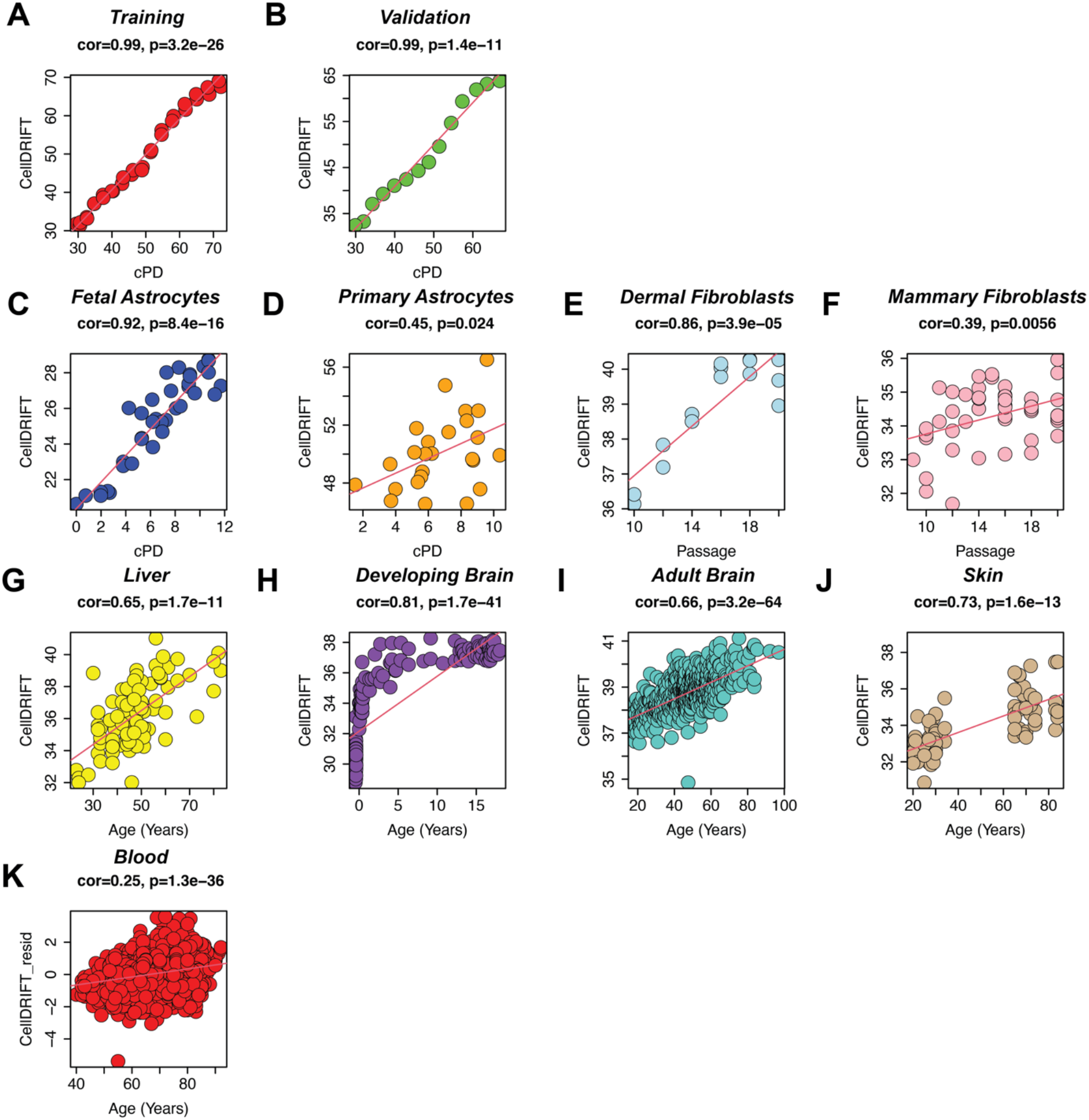
Training and validation of CellDRIFT. **(A)** Training and **(B)** Validation of PC measure CellDRIFT, trained from yellow and tan module CpGs (2,322 total) and hTERT immortalized replication data. More training information is available in the Methods. *In vitro* validation **(C-F)** and *in vivo* multi-tissue validation **(G-K)**. Note, blood was residualized by cellular composition (b-cells, granulocytes, CD8T cells and monocytes). Age correlations and statistical significance was determined via Pearson correlations.

**Supplemental Figure 7:**
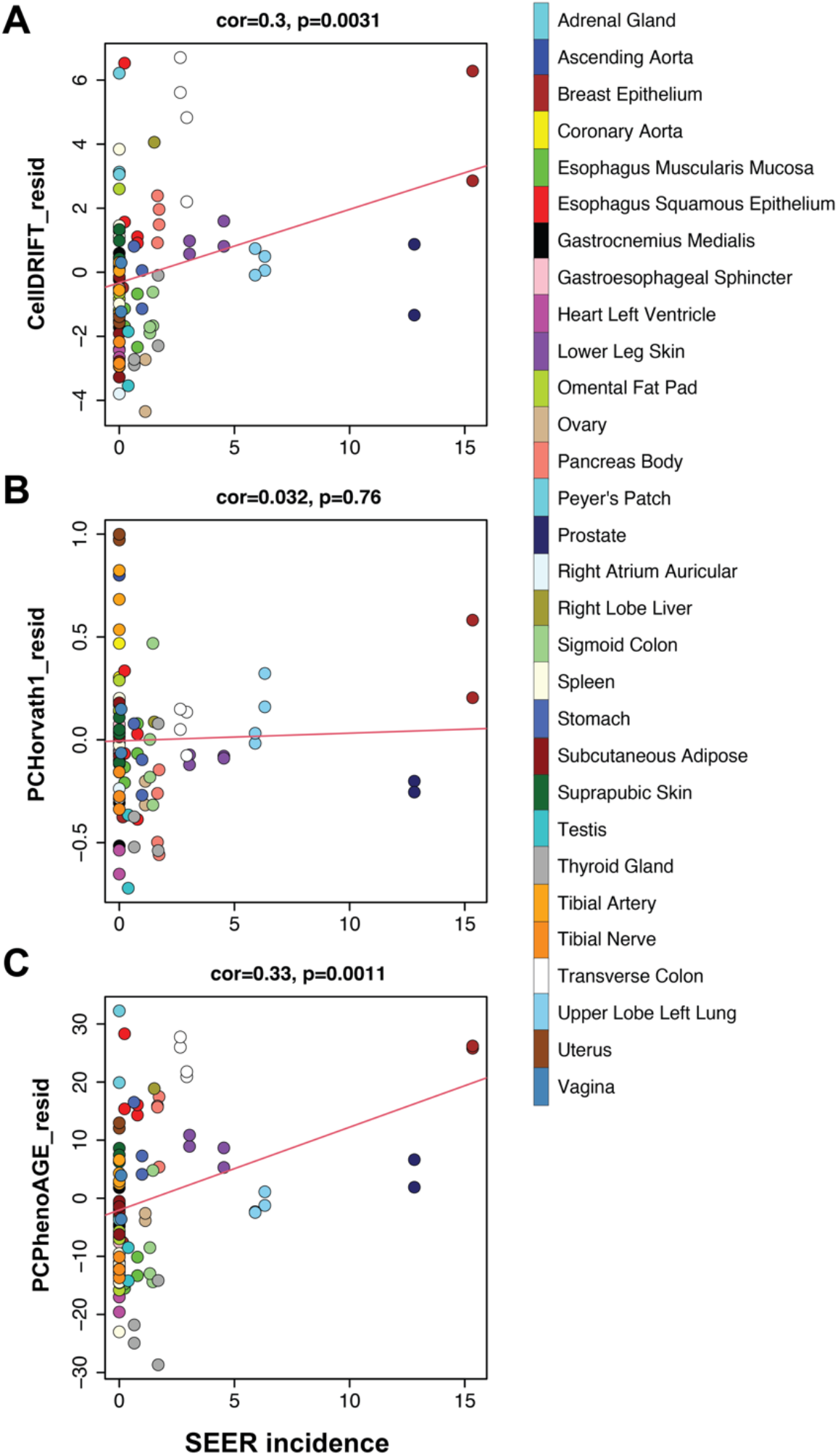
Complete whole-body tissue dataset inclusive of near zero risk cancer tissues. DNAmAge analysis of **(A)** CellDRIFT**, (B)** PCHorvath1 and **(C)** PCPhenoAGE in relation to SEER incidence of 29 tissues from 4 healthy donors. All DNAmAge scores were residualized by age.

**Supplemental Figure 8:**
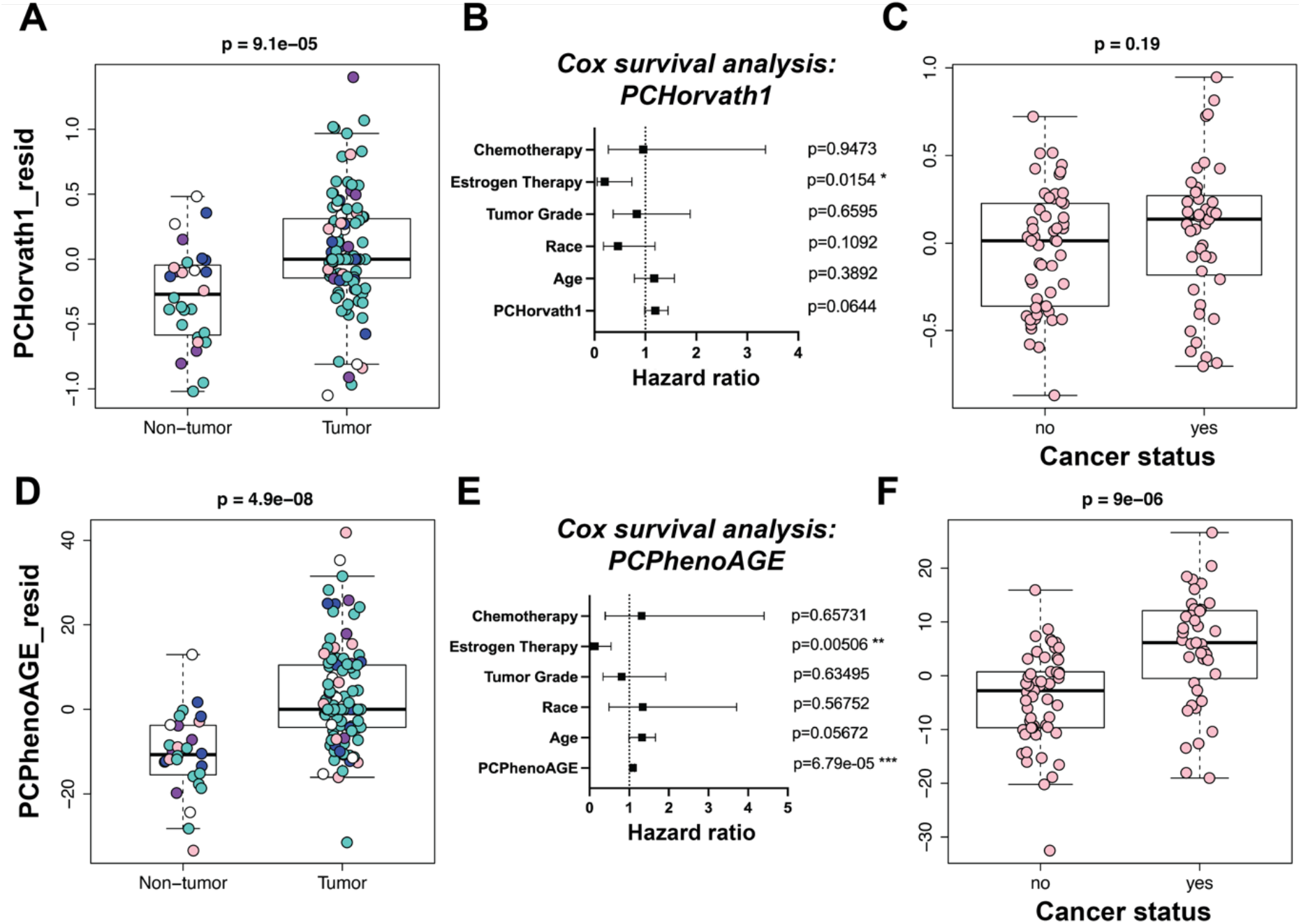
Cancer detection using classically trained *ex vivo* clocks. Plots displaying cancer findings of PCHorvath1 and PCPhenoAGE measures. **(A,D)** Pooled cancer detection of breast, colon, lung, thyroid and pancreas cancer samples from GSE53051. Pink=Breast, Teal=Thyroid, White=Lung, Blue=Colon, Purple=Pancreas. DNAmAge scores were residualized by age, sex and tissue type. **(B,E)** COX survival analysis of breast cancer patients from GSE37754. Note, the hazard ratio for Age and PCclocks was calculated as a risk increase per 10 years of life. **(C,F)** Differences in DNAmAge in healthy breast tissue of known breast cancer patients and participants with no history of prior breast cancer (Rozenblit et. al 2022) [37]. DNAmAge scores were residualized by age prior to analysis.

## Methods

### Experimental

#### Fetal astrocyte extraction

4 fetal-mortal astrocyte cell lines were derived from the cerebral cortex of two different donors (ScienCell #1800). Donor 1, which was split into -hTERT1, -hTERT2 and -hTERT3 cell lines and was used in training experiments and Donor2 which was split into -hTERT4. Tissue was received by ScienCell Research Laboratories from non-profit tissue providers, obtained with informed consent of donor’s family aged over eighteen, and under established protocols in compliance with an institutional review board and local, state, and federal laws. No payment, commercial rights or financial rights were provided to the donor family. Further details can be obtained from ScienCell Research Laboratories.

#### Primary astrocyte extraction

3 primary human astrocyte cell lines were derived from the cerebral cortex of one 21-year-old male donor (Creative Biolabs, #NCL-2103-P104). The 21M donor was split into Astro1, Astro2 and Astro3 cell lines, which were then subsequently exhaustively passaged. All tissue collection procedures used by Creative Biolabs, and its partners, was performed in compliance with institutional review boards and local, state, and federal laws. Further details can be obtained from Creative Biolabs.

#### Immortalized (hTERT) astrocyte preparation

hTERT immortalized fetal astrocyte cell lines were supplied by Applied biological materials (Abm) (#T0281). The immortalized cells were supplied at passage 12, with passage reporting starting after hTERT immortalization. Upon receiving, we subsequently split the immortalized fetal donor into 3 hTERT cell lines, +hTERT1, +hTERT2 and +hTERT3, which were then passaged to p27. Further details about plasmids used in transfection and details about donor sourcing can be obtained from Abm.

#### Astrocyte replicative passaging and cellular culturing

Fetal cells were exhaustively passaged and split a total of 10 times (9-15 cumulative population doublings, depending on replicate), where β-gal activity (C12FDG) was measured using flow cytometry or confocal microscopy at each passage to confirm exhaustive replication was achieved. Primary astrocytes were also exhaustively passaged and split a total of 10 times (8-9 cumulative population doublings, depending on replicate). hTERT immortalized astrocytes were passaged a total of 27 times (73-75 cumulative population doublings, depending on replicate).

All cell lines were seeded at 8,000 cells/cm^2^ (0.5×10^6^ cell/p100) with appropriate growth media and supplements (complete astrocyte medium containing amino acids, vitamins, hormones, trace minerals, 2% fetal bovine serum and 1% PEN/STREP in HEPES pH 7.4 bicarbonate buffer, ScienCell #1801) to promote cell adhesion and growth. Of note, Poly-L-Lysine was not required for adequate cell adhesion. All cells were grown under normoxic conditions (20% O_2_, 5% CO_2_) at 37°C. Cells were split (0.05% Trypsin-EDTA (Gibco, #25300-54)) when they reached approximately 90% confluence or when static growth was achieved. Cells were counted using the Invitrogen countess and cell counting chamber slide with trypan blue. Cumulative population doubling was calculated using the initial and final cell density, as determined by the countess (2^x^=FD/ID, where x=population doubling, FD=final cell density and ID=initial cell density).

#### DNA preparation

Longitudinal samples were collected at every passage and DNA was extracted using the Qiagen DNAeasy Blood and Tissue Kit (#69504). Note, samples were treated with proteinase K and RNAse A and eluted with 200 μl elution buffer. Following final elution, DNA was verified using nanodrop (Thermo Scientific) and Qubit fluorometer (Invitrogen). Spin concentration was used as necessary with low DNA content samples. Prior to library preparation we used a Qubit fluorometer (Invitrogen) to quantify the extracted genomic DNA.

#### Beta-galactosidase confocal microscopy

To assess senescence status, we employed a beta-galactosidase imaging method. In brief, cells were split into 12-well dishes (0.125×10^6^ cells/well) with a glass cover slide at the bottom of each well and allowed to settle for 24 hours. Cells were first pre-treated with Bafilomycin A1 (Selleckchem: S1413, 622.83 g/mol, 100 μM stock). Existing media was aspirated, then cells were washed with PBS and replaced with treated Bafilomycin A1 media for 30 min at a final concentration of 100 nM. Following Bafilomycin A1 pretreatment to normalize lysosome activity, C12FDG (Invitrogen, #D2893, 853.92 g/mol, 10 mM stock) was added directly to the existing media for 90 min at a final concentration of 10 μM. Note, due to light sensitivity, exchange was conducted in a dark environment. Following Bafilomycin A1 and C12FDG treatment, media was aspirated, and cells were washed with PBS 3x, fixed with 4% PFA/PBS (10 min), followed by 2x PBS washes and then counter stained with DAPI (Invitrogen, #P36935) and mounted onto coverslips. Fixed cells were immediately imaged using a ZOE fluorescent cell imager (Bio Rad). Percent cell positively was calculated using ImageJ with a background/negative threshold value. Any cells above this threshold were considered positive. All cells in each image frame were counted.

#### F-actin (Phalloidin-CruzFluor 532) confocal microscopy

Passage 3, 6 and 10 fetal astrocytes (-hTERT1-3) were concurrently passaged to dynamically visualize changes in cellular morphology. In brief, cells were split into 12-well dishes (0.125×10^6^ cells/well) with a glass cover slide at the bottom of each well. Cells were allowed to grow for 5 days before assessment. Following growth, cells were fixed using 4% formaldehyde in PBS for 30 minutes. In summary, loose cells and media were aspirated, then washed 2x with PBS, then fixed with 4% formaldehyde for 30 minutes. After fixation, cells were washed 2x with PBS, then Phalloidin-CruzFluor 532 (1x) (Santa Cruz, #363793) conjugate was added for 90 minutes. Following conjugation, cells were gently rinsed 2x with PBS, then 2x with H20. Finally, cells were counter stained with DAPI (Invitrogen, #P36935), mounted onto coverslips and imaged with a ZOE fluorescent cell imager (Bio Rad).

#### Absolute telomere length qPCR quantification

Telomere attrition was calculated as a percent change from baseline using absolute telomere length assessed using a qPCR kit from ScienCell (#8918). -hTERT1 and hTERT1 cell lines were used in the assessment. More specifically, -hTERT cells passaged 2, 4, 6, 8 and 10x and hTERT1 cells passaged 13, 15, 17, 19, 21, 23, 25, 27x were used in the final telomere attrition reporting and comparisons.

### Computational

#### Data processing

All samples were assigned a single-blinded code and randomized for library preparation and sequencing to control for any batch errors. DNAm data was generated using the Infinium HumanMethylation850 BeadChip and preprocessed using minfi45 [38] and normalized using the noob method56 [39]. Prior to analysis all sex chromosome CpGs were excluded, and for training and validation purposes we aligned the CpGs to the 450k array, giving a final matrix of 442,242 CpGs.

#### Training and validation of DNAmImmort, Module clocks, CellDRIFT and PC clocks

R was the primary platform used throughout the study (Version 4.1.1). Prism (Version 9) was also used for certain statistical analysis and plotting. DNAmImmort was constructed using only passaged hTERT cells. In brief, 3 hTERT cell lines were used in the training and validation process, all from the same donor and immortalization pair. +hTERT1 and +hTERT2 (n=31, p13-p27) was used in training, hereafter referred to as hTERT training, and +hTERT3 (n=14, p13-p27) was used in validation, hereafter referred to as hTERT validation. In summary, PCA was conducted on the hTERT training samples, then we used elastic net regression modeling to select the final 16 PCs used in the measure based on using cPD as the training variable for calibration. The 16 PCs were comprised of PCloading scores for all 442,242 CpGs, with elastic net coefficients based on lambda penalty values representing the lowest mean-squared error, selected via 10-fold cross validation.

From modules identified according to WGCNA in the following section, we trained a number of smaller clocks. These were trained using the same hTERT training and validation samples, except PCA was conducted on the module CpGs, resulting in a measure with significantly reduced CpGs (Turquoise=4,008 CpGs, Blue=3,859 CpGs, Brown=2,641 CpGs, Yellow=2,025 CpGs, Green=1,525 CpGs, Red=1,252 CpGs, Black=912 CpGs, Pink=882 CpGs, Magenta=880 CpGs, Purple=756 CpGs, Greenyellow=721 CpGs, Tan=297 CpGs, Salmon=234 CpGs, Cyan=108 CpGs). See more details below on the process of isolating the most physiologically relevant module CpGs.

To determine the cell and tissue age correlations we first conducted PCA in each module CpG cohort (eigengene) in our *in vitro* replication data and then calculated the resulting variance (PC1) within each validation dataset (Sup Fig. 4B). Next, we trained elastic net selected PC clocks (DNAmModuleColor) as replication predictors in our *in vitro* model and then calculated DNAmAge Pearson correlations within each module clock and validation dataset (Fig. 1D). In both eigengene (PC1) and module clock analysis, the yellow and tan modules CpGs were the most physiologically relevant replication fingerprints, with strong correlations across both *in vivo* tissues - liver, skin, developing and adult brain, and blood – and *in vitro* cell lines – fetal astrocytes, primary astrocytes, primary dermal fibroblasts and primary mammary fibroblasts (Fig. 1D). In moving forward, the CpGs in these two modules served as the input CpGs for the final composite measure, CellDRIFT.

The final cellular replication measure - CellDRIFT (**<u>Cell</u>**ular **<u>D</u>**ivision and **<u>R</u>**eplication **<u>I</u>**nduced **<u>F</u>**ingerprin**<u>T</u>**) - was trained from the combined CpGs from the yellow (2,025) and tan (297) modules, which were selected from extensive *in vitro* and *in vivo* validation analysis to determine multi-tissue and physiological relevance (Fig. 1D). Instead of recalculating PCs for the new 2,322 CpG cohort, we combined the independent PCs (62 total, 31 from each module) and conducted elastic net to select for the final PCs for inclusion in the final measure. This way, each signal had equal weight and possibility for contributing to the final measure and CpG occupancy wasn’t a biasing factor. The following PCs were selected from each module: Yellow [PC1, PC2, PC3, PC10, PC17] and Tan [PC1, PC2, PC3, PC4, PC5, PC7, PC10, PC11]. Further details on PC-trained measures can be found in our previous reports [5,11]. A package is under construction for easy calculation of CellDRIFT from external datasets. See attached Data Supplement 1 for a list of the CpG sites in each module.

**Data Figure 1:**
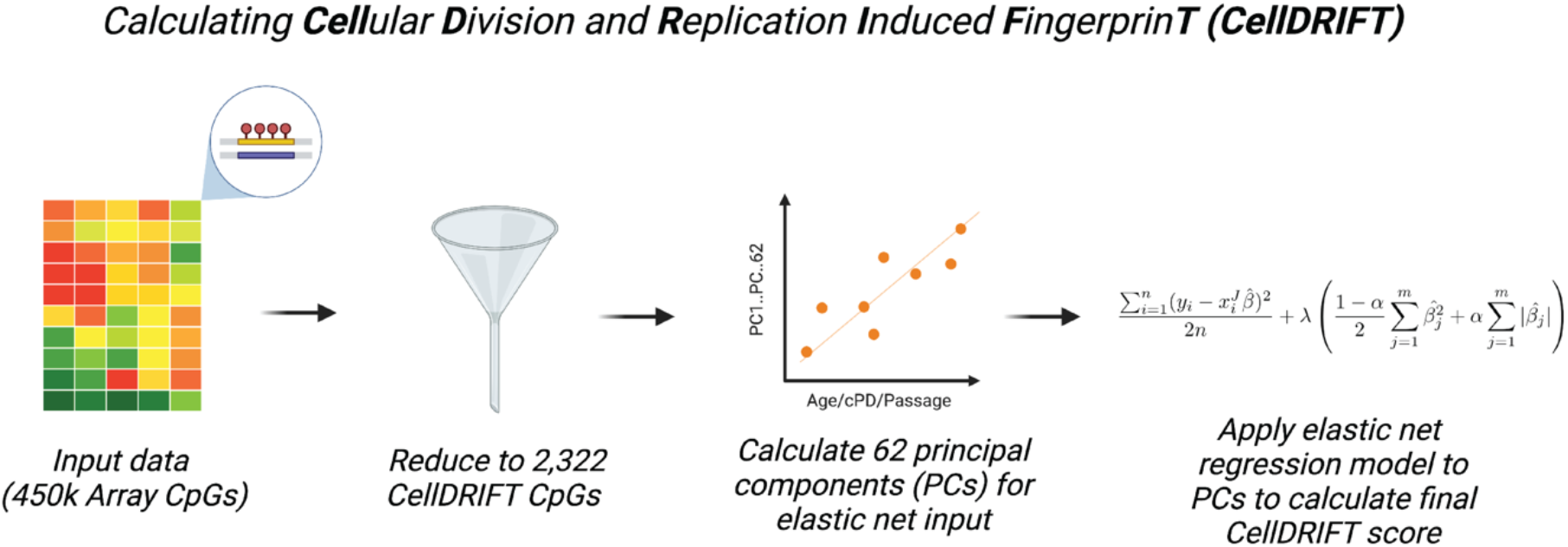
CellDRIFT calculation pipeline. Data schematic showing process of selecting 2,322 CellDRIFT CpGs, calculating the 62 PCs, then feeding them into the elastic net regression resulting in the final CellDRIFT score.

Validation of all measures was done first by assessing the DNAmAge of the hTERT validation data (n=14), followed by various multi-tissue and additional *in vitro* datasets (Sup Fig. 3). Pearson correlations were used to determine associations between all measures and validation data presented in Fig. 1D. Note, liver, developing brain, adult brain, blood and skin were assessed as age-correlations, hTERT, fetal astrocytes and primary astrocytes were assessed as cumulative population doublings (cPD) correlations, and dermal and mammary cell lines were assessed as passage correlations due to limited cPD data information.

As controls, we evaluated the traditionally trained *ex vivo* clocks PCHorvath1 and PCPhenoAGE and found CellDRIFT outperformed the multi-tissue measure PCHorvath1, and performed similarly to PCPhenoAGE, which is remarkable considering PhenoAGE was trained as a health span predictor and is highly associated with mortality, while CellDRIFT is only trained from *in vitro* cell divisions (Sup Fig. 8). We associated the improved cancer detection power of PCPhenoAGE over PCHorvath1 to its tighter association with all-cause mortality. PC clock calculations for PCHorvath1 and PCPhenoAge were conducted using the data analysis pipeline from Higgins-Chen et. al 2022 [13].

##### WGCNA module construction and hierarchical clustering

We conducted consensus WGCNA [20] and hierarchical clustering to produce distinct DNAmAge signals, as we previously reported [7]. In brief, we used 2 input datasets (hTERT training and liver aging), with the remaining datasets (hTERT validation, blood, brain, skin, primary astrocyte, fetal astrocyte, dermal and mammary fibroblasts) excluded for validation purposes. In total, we used 20,101 CpGs in the consensus analysis. The CpGs included were the top driver CpGs of DNAmImmort, which were selected from normalizing PC loading scores and selecting the top values (Sup Fig. 2F). In brief, adjacencies were estimated for each dataset, which is based on biweight midcorrelations. These adjacencies were then converted to Topological Overlap Matrices (TOMs) where a minimum dissimilarity score was calculated for each CpG pair across the two TOMs. Hierarchical clustering was then conducted with the following parameters, deepSplit= 1, cutHeight = 0.95, minClusterSize = 50, and distance = 1-consensus TOM, method = average. The resulting network produced 14 modules. No further module cutting was preformed, with all CpGs in the input analysis being assigned a module. Following module construction, we estimated PC1 for each module using the hTERT training data (Sup Fig. 4B) and then applied this score to all validation data to serve as a validation metric of module connectivity and similarity. Module clocks were then trained from the module CpGs and taken forward to determine differences in DNAmAge between modules. For further information on module development refer to Minteer et. al 2022 [7].

##### Cistrome genome enrichment analysis

In order to better determine the functionality of the CpGs selected from each module we used the Cistrome gene analysis tool kit (http://dbtoolkit.cistrome.org/) to determine enriched genes and histone marks in each CpG dataset. In brief, we plotted all transcription factors and chromatin regulators with significant overlap with the CpG modules and then plotted a heatmap of the top 5 genes (average Giggle score across all GSM_ID hits) from each module together, in order to compare module to module enrichment (Sup Fig. 5). Note, the summary module enrichment analysis used the top 100 CpGs from each module, determined from the kME score. Enriched genes were normalized by selecting 100 background CpGs from the original 440k training dataset and correcting for each GSM_IDs Giggle score. Enrichment analysis is displaying the average Giggle score across all GSM_IDs, with the top 5 for each module plotted in Sup Fig. 5Q and then the top 20 were fed into String analysis in Fig. 2B. Note, histone mark analysis was conducted on the top 100 kME CpGs and corrected for background hits with the final heatmap displaying the top 5 marks by average Giggle score (Fig. 2A). Giggle score is a rank of genome significance between genomic locations of query file and thousands of genome files from databases like ENCODE. For further information on normalization and plotting Giggle scores refer to Minteer et. al 2022 [7].

##### Additional analysis pipelines

Genomic partitioning and CpG locations were determined using LolaWeb (http://lolaweb.databio.org/). STRING protein-protein network analysis was conducted using the STRING database (https://string-db.org/). Medium confidence (0.4) was set as the minimum interaction score threshold.

##### Statistical analysis and R packages

Plotting and module development were conducted using the WGCNA package. Additional plotting was done using the ggplot2 package. Elastic net modeling was conducted using the glmnet package. Survival analysis and COX hazard ratio analysis was conducted using the survival, survminer and dplyr packages. Hazard ratios were calculated in reference to survival data from a cohort of breast cancer patients (GSE37754) with the interaction variable being the CellDRIFT or PCclock scores. Additional packages of interest used in this study were: BiocManager, lattice, viridis, RColorBrewer, reshape, and GEOquery. Pearson correlations were used to assess age and passage associations, the Kruskal–Wallis ONE-way ANOVA test were used for multi-group comparisons, Two-tailed t tests were used to compare group-group significance and biweight midcorrelations were used for determining adjacencies in module construction analysis.

##### Data accessibility and usage

All data used in this study is summarized below: *<u>Training</u>:* hTERT training data (this study, n=31 samples), *<u>Validation</u>:* hTERT validation data (this study, n=14), Liver aging (GSE48325), Brain aging (GSE74193), Blood aging (GSE40279), Skin aging (GSE52980), Primary Astrocytes (this study, n=25), Fetal Astrocytes (this study/GSE202554), Primary Dermal Fibroblast and Primary Mammary Fibroblasts (E-MTAB-8327), *<u>Cancer</u>:* Pooled cancer samples (GSE53051), Breast cancer survival data (GSE37754), Healthy breast tissue - Rozenblit et. al 2022 [37], Whole-body tissue (ENTEx study), *<u>Reprogramming</u>:* Yamanaka fibroblast re-programming and passaging data (GSE54848) and iPSC/ESC passaging data (GSE31848).

## Acknowledgements

Contribution is based on authorship. We thank our commercial and academic partners for access to data for validating our study.

## Author Contributions

Conceptualization: CJM, MEL

Methodology: CJM, MEL, KT, PN, JR, JL, MF, TC, EH, LP, MG, MR, KB

Investigation: CJM, MEL

Visualization: CJM, MEL

Funding acquisition: MEL

Project administration: MEL

Supervision: MEL

Writing – original draft: CJM

Writing – review & editing: CJM, MEL, KT, PN, JR, JL, MF, TC, EH, LP, MG, MR, KB

## Data and materials availability

All data used in the study is listed in the supplementary material. Any data not public can be requested.

